# Sparse CBX2 nucleates many Polycomb proteins to promote facultative heterochromatinization of Polycomb target genes

**DOI:** 10.1101/2024.02.05.578969

**Authors:** Steven Ingersoll, Abby Trouth, Xinlong Luo, Axel Espinoza, Joey Wen, Joseph Tucker, Kalkidan Astatike, Christopher J. Phiel, Tatiana G. Kutateladze, Tao P. Wu, Srinivas Ramachandran, Xiaojun Ren

**Affiliations:** Department of Chemistry, University of Colorado Denver, Denver, CO 80217-3364, USA; Department of Biochemistry and Molecular Genetics, University of Colorado School of Medicine, Aurora, CO, USA; Department of Molecular and Human Genetics, Baylor College of Medicine, Houston, TX, USA; Department of Integrative Biology, University of Colorado Denver, CO 80217-3364, USA; RNA Bioscience Initiative, University of Colorado School of Medicine, Aurora, CO, USA; Department of Pharmacology, University of Colorado School of Medicine, Aurora, CO 80045, USA

**Keywords:** Polycomb, PRC1, PRC2, CBX2, H3K27me3, chromatin, nucleation, liquid-liquid phase separation, facultative heterochromatin, constitutive heterochromatin, single-molecule imaging, live-cell single-molecule tracking

## Abstract

Facultative heterochromatinization of genomic regulators by Polycomb repressive complex (PRC) 1 and 2 is essential in development and differentiation; however, the underlying molecular mechanisms remain obscure. Using genetic engineering, molecular approaches, and live-cell single-molecule imaging, we quantify the number of proteins within condensates formed through liquid-liquid phase separation (LLPS) and find that in mouse embryonic stem cells (mESCs), approximately 3 CBX2 proteins nucleate many PRC1 and PRC2 subunits to form one non-stoichiometric condensate. We demonstrate that sparse CBX2 prevents Polycomb proteins from migrating to constitutive heterochromatin, demarcates the spatial boundaries of facultative heterochromatin, controls the deposition of H3K27me3, regulates transcription, and impacts cellular differentiation. Furthermore, we show that LLPS of CBX2 is required for the demarcation and deposition of H3K27me3 and is essential for cellular differentiation. Our findings uncover new functional roles of LLPS in the formation of facultative heterochromatin and unravel a new mechanism by which low-abundant proteins nucleate many other proteins to form compartments that enable them to execute their functions.

## INTRODUCTION

Facultative heterochromatinization of developmental master regulators is essential for the generation and maintenance of diverse cell types in metazoans ^1–3^. Dysregulation of facultative heterochromatin is associated with many diseases, such as cancer and developmental disorders ^4–6^. A hallmark of facultative heterochromatin is the presence of Polycomb group (PcG) proteins and their enzymatic product H3K27me3 ^1–3^. Facultative heterochromatin forms compartments that are separated from compartments of constitutive heterochromatin which features DNA-dense regions marked by H3K9me3 ^7–9^. There is a paucity of information concerning molecular factors that demarcate the spatial boundaries of facultative heterochromatin and constitutive heterochromatin.

Polycomb repressive complex 2 (PRC2), one of the PcG complexes, functions as a H3K27-specific methyltransferase and generates H3K27me3 which covers broad regions of chromatin termed Polycomb domains ^10–12^. PRC2 deposits H3K27me3 through a two-step mechanism: PRC2 is recruited to a limited number of spatially interacting nucleation sites and then spreads H3K27me3 to distal regions through long-range contacts ^13–19^. PRC2 is engaged with nucleation sites through its accessory subunits that directly interact with chromatin ^15,20–22^, stabilizing PRC2 at chromatin ^23–26^ and facilitating trimethylation of H3K27 ^25,26^. Both the allosteric activation of PRC2 by H3K27me3 and spatial clustering of Polycomb targets enable the long-range spreading of H3K27me3 ^15,18^. However, the factors which spatially cluster Polycomb targets for the efficient deposition of H3K27me3 by PRC2 remain poorly understood.

Currently, a widely held presumption is that proteins with very low abundance are considered less functionally relevant, which is partly due to limitations in detecting and quantifying them. However, low-abundant proteins can play critical roles in nucleation, assembly, and amplification of condensation ^27–30^. This behavior is recently exemplified by the long non-coding RNA Xist ^29,30^. There are ∼200 copies of Xist within an individual cell, which forms ∼50 diffraction-limited foci ^31,32^. Xist foci have ∼2 RNA molecules, which nucleate protein complexes to transcriptionally silence one X chromosome. The expression of CBX2, a component of the canonical form of PRC1 (cPRC1, also termed CBX-PRC1), is very low in mouse embryonic stem cells (mESCs) ^33–35^. CBX2 is the major driver of the formation of CBX-PRC1 condensates ^36–39^ and is important for proper development and differentiation ^40–43^. Due to the low abundance of CBX2 in mESCs, it is unclear whether sparse CBX2 plays a functional role in mESCs.

CBX-PRC1 has been shown to form condensates *in vitro* and in cells ^39,44–48^. These condensates are involved in repressing Polycomb targets ^41,43,46^. It was thought that each of the CBX-PRC1 complexes consists of one CBX protein (CBX2/4/6/7/8), one RING1A or RING1B, one MEL18 or BMI1, and one PHC protein (PHC1/2/3) ^49,50^; however, the stoichiometry of CBX-PRC1 proteins within condensates remains unknown. The emerging model is that CBX-PRC1 condensates are multi-component assemblies, and that CBX proteins drive condensate formation ^36,38,39,47^. The assembly of condensates is modulated by the composition and concentration of CBX-PRC1 proteins ^39,47^. Additionally, the PHC family of proteins can polymerize and form condensates and also play an important role in the repression of Polycomb targets ^46,51–54^. The combination of the polymerization ability of PHC and the condensation ability of CBX confers contextually specific biophysical properties of CBX-PRC1 condensates ^47^. Furthermore, the expression pattern of Polycomb paralogs changes during cell differentiation and is distinct in different cell types ^10,35^, therefore, it is important to develop the approaches that would enable quantification of the molecular composition and stoichiometry of Polycomb proteins within condensates in specific cell types.

Similar to PRC1, there are four core subunits of PRC2: EZH2 or EZH1, SUZ12, EED, and RbAp46/48 ^13,14^. At the molecular level, H3K27me3, the product of the PRC2 catalytic activity, provides a docking site for the CBX-PRC1 complexes ^11,12^, and thus PRC2 and CBX-PRC1 coordinate their functions to establish and maintain Polycomb domains. PRC2 has been shown to appear as condensates within cells ^23,55,56^. However, it remains unclear whether there is a physical and functional interplay between CBX-PRC1 condensates and PRC2 condensates.

Membrane-less condensates formed by phase separation have emerged as a promising principle for organizing nucleic acids and proteins within the cell ^57–67^. Phase separation has been linked to proteins and nucleic acids that are involved in active transcription ^68–79^, chromatin organization ^80–83^, and facultative and constitutive heterochromatin formation ^36,38,56,84–89^. Phase-separated condensates can concentrate proteins and nucleic acids, which provide the molecular principles by which liquid-liquid phase separation (LLPS) accelerates biochemical reactions ^57,63,65^. In fact, recent studies have demonstrated that LLPS accelerates the target search of Polycomb CBX2 ^37^. Therefore, we speculate that sparse CBX2 could seed the formation of Polycomb condensates that control the spatial boundaries of facultative heterochromatin and enhance the efficiency of H3K27me3 deposition. Here, using genetic engineering, molecular approaches, and single-molecule cell imaging, we determine that ∼3 CBX2 proteins nucleate the formation of non-stoichiometric Polycomb condensates, which leads to the enrichment of ∼70 RING1B and ∼86 EZH2 per condensate in mESCs. The LLPS capacity of sparse CBX2 regulates the spatial boundaries of facultative heterochromatin and modulates the deposition of H3K27me3, which is essential for cellular differentiation. Together, our results indicate that low-abundant proteins can function to nucleate many proteins, which form large assemblies through phase separation.

## RESULTS

### ∼3 CBX2 proteins per one non-stoichiometric Polycomb condensate

The molecular stoichiometry of phase-separated condensates in living cells remains unclear, yet stoichiometry is essential for understanding condensate assembly and function. Here, we developed a generic approach that enables quantifying the molecular stoichiometry of phase-separated condensates by combining CRISPR-Cas9-mediated genetic engineering, quantitative immunoblotting, and quantitative single-molecule sensitive fluorescence microscopy. Through CRISPR-Cas9-mediated genetic engineering ^90^, we inserted HaloTag to the N- or C-terminus of the endogenous locus of PRC1 genes (*Cbx2*, *Cbx7*, *Ring1a*, *Ring1b*, *Mel18*, and *Phc1*) and a PRC2 gene (*Ezh2*) in mESCs (**Figure 1A** and **S1A-B**). Following the labelling of HaloTag with TMR ligand, we performed live-cell fluorescence microscopy imaging. Since CBX2 was undetectable when using confocal and epifluorescence microscopy without a single-molecule sensitivity, we took micrographs of cells by using single-molecule HILO microscopy (**Figure 2F**). All tested proteins of the PRC1 and PRC2 complexes were enriched within condensates. We first quantified the condensed fraction (the fraction of proteins within condensates vs total proteins) of each labeled Polycomb subunit (**Figure 1B**). CBX2 had the highest condensed fraction (∼12%) while EZH2 had the lowest (∼3%). Other PRC1 proteins had condensed fractions ranging from ∼ 4% to ∼8%. The number of condensates also varied between Polycomb proteins. CBX2 had the most condensates per cell (∼100) while EZH2 had the fewest (∼44) (**Figure 1C**). The condensed fraction and the number of CBX2 condensates were similar among three distinct cell lines (**Figure S1A**), which is consistent with previous reports ^39^. These results indicate that both PRC1 and PRC2 are enriched within condensates in mESCs but to a different degree.

**Figure 1.**
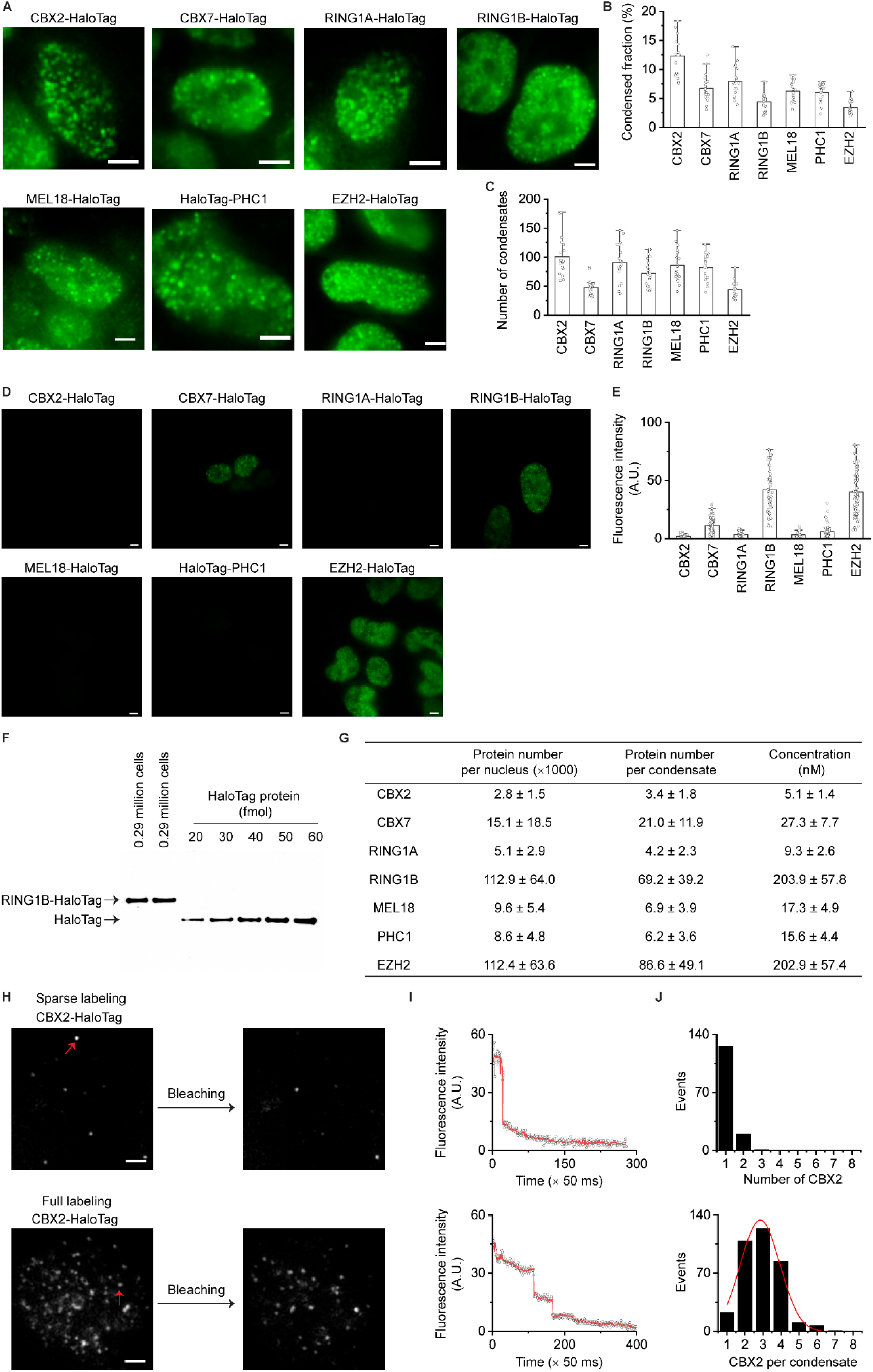
∼3 CBX2 proteins per condensate in mESCs. (**A**) Live-cell epifluorescence images of HaloTag Polycomb fusion proteins. HaloTag was inserted into the N- or C-terminus of genetic loci of PRC1 subunits (CBX2, CBX7, RING1A, RING1B, MEL18, and PHC1) and the C-terminus of genetic loci of PRC2 subunit (EZH2) in mESCs. Micrographs were taken by using single-molecule sensitive highly inclined thin illumination (HILO) microscopy (Figure 2F) since CBX2-HaloTag was undetectable when using fluorescence microscopy without a single-molecule sensitivity (unless otherwise indicated, all epifluorescence images were taken by single-molecule HILO microscopy). To show the distribution of proteins, images were taken and presented by using different settings. Scale bar, 5.0 µm. (**B**) Bar overlap plot of the condensed fraction of HaloTag Polycomb fusion proteins quantified from Figure 1A. Condensed fraction is defined as the fraction of condensed proteins to total proteins within the nucleus. (**C**) Bar overlap plot of the number of condensates of HaloTag Polycomb fusion proteins quantified from Figure 1A. (**D**) Live-cell epifluorescence images of HaloTag Polycomb fusion proteins. To compare fluorescence intensities, images were taken and presented by using the same settings. Scale bar, 5.0 µm. (**E**) Bar overlap plot of the fluorescence intensity of HaloTag Polycomb fusion proteins quantified from Figure 1D. RING1B-HaloTag, MEL18-HaloTag, PHC1-HaloTag, and EZH2-HaloTag are heterozygous while others are homozygous. The final intensity was normalized as homozygosity. (**F**) Immunoblotting analysis of the RING1B-HaloTag protein extracted from mESCs and recombinant HaloTag protein with a series of dilutions. Blot was probed by using anti-HaloTag antibodies. (**G**) Estimated protein number per nucleus, protein number per condensate, and concentration for HaloTag Polycomb fusion proteins. Error represents standard deviation from three biological triplicates. (**H**) Single-molecule photobleaching of CBX2-HaloTag sparely (top) or densely fully (bottom) labelled with HaloTag ligand JF549. Cells were fixed by 4.0% paraformaldehyde and imaged at single-molecule conditions. Red arrowheads indicate fluorescent spots/condensates to be bleached. Scale bar, 5.0 µm. (**I**) Representative photobleaching curve. The photobleaching steps were detected by Chung-Kennedy filter (red line). The number of individual steps represents the number of proteins within a spot. (**J**) Histogram of single-molecule photo-bleaching steps of CBX2-HaloTag that were sparsely (top) and fully (bottom) labelled, respectively. The bottom histogram was fitted by a Gaussian function to estimate average protein number per condensate.

**Figure 2.**
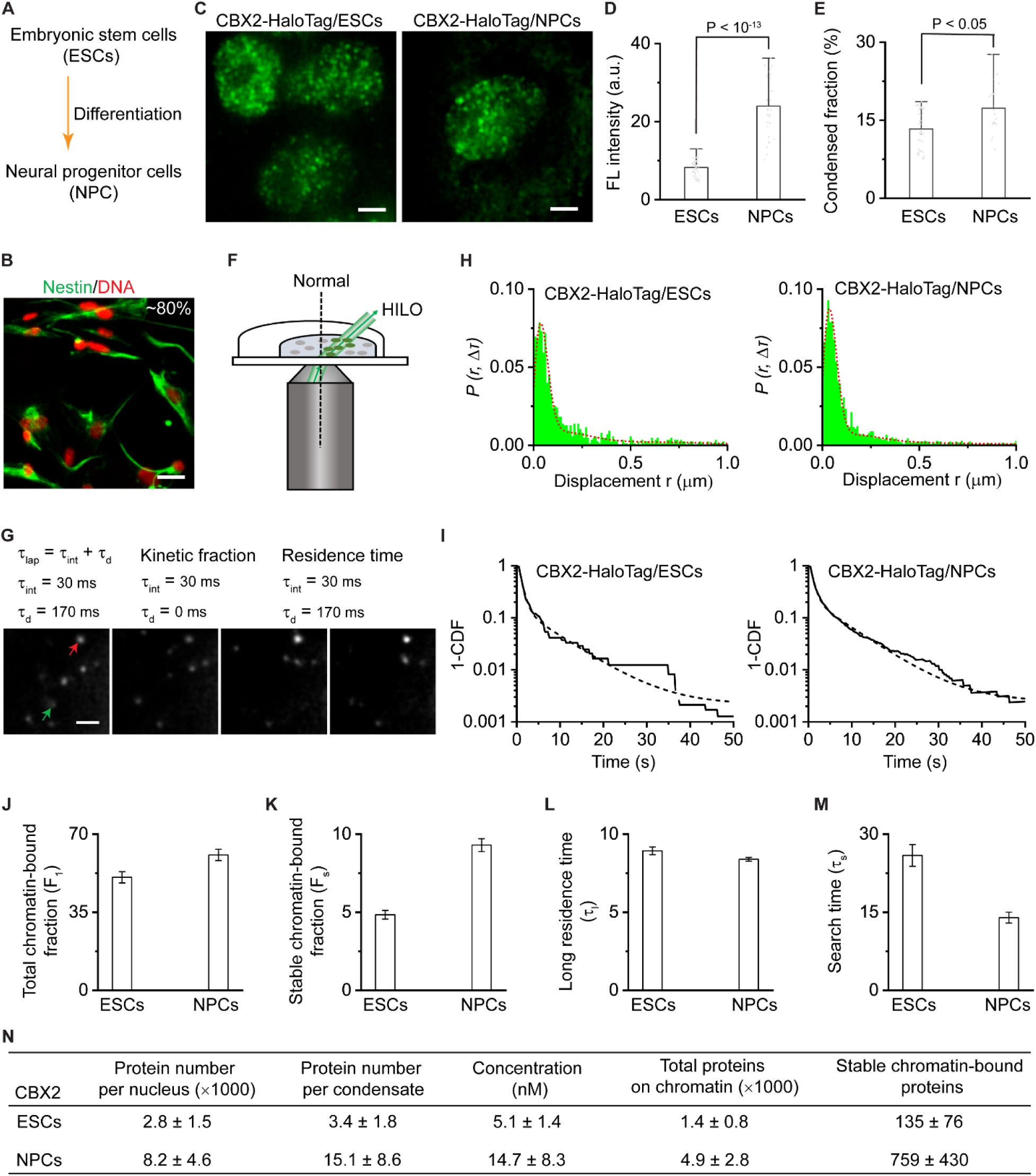
Cell-type-dependent control of the number of CBX2 proteins per condensate and of the interacting of CBX2 with chromatin. (**A**) Overview of experimental approaches. mESCs were differentiated into neural progenitor cells (NPCs). (**B**) Immunostaining of mESCs-derived NPCs by anti-Nestin antibodies. DNA was stained using Hoechst. A representative overlay image is shown. Nestin is green and Hoechst is red. Scale bar, 5.0 µm. (**C**) Live-cell epifluorescence images of CBX2-HaloTag in mESCs and NPCs. To show distributions, images were taken under different settings. Scale bar, 5.0 µm. (**D**) Bar overlap plot of the fluorescence intensity of CBX2-HaloTag in mESCs and NPCs. Images used for quantification were taken under the same settings. P value, student’s t-test with two-tailed distribution and two-sample unequal variance. (**E**) Bar overlap plot of the condensed fraction of CBX2-HaloTag in mESCs and NPCs. P value, student’s t-test with two-tailed distribution and two-sample unequal variance. (**F**) Schematic representation of single-molecule HILO microscopy used for live-cell single-molecule tracking (SMT). (**G**) Example images showing individual CBX2-HaloTag molecules. The red arrowhead indicates that a CBX2-Halotag molecule stays at chromatin and the green arrowhead represents that a CBX2-HaloTag molecule freely diffuses. For studies of kinetic fraction, the camera integration time is 30 ms with no dark time between camera exposures. For estimating of residence time, the integration time is 30 ms and the dark time is 170 ms. Scale bar is 5.0 µm. (**H**) Displacement histogram for CBX2-HaloTag in mESCs (N = 22 cells, n = 3082 displacements) and NPCs (N = 33 cells, n = 10666 displacements). The short dash red curve indicates the overall fit. (**I**) Survival probability distribution of the dwell times for CBX2-HaloTag in mESCs and NPCs. The distributions were fitted with a two-component exponential decay model (dotted lines). (**J-K**) Total chromatin-bound fraction (F_1_) and stable chromatin-bound fraction (F_s_) for CBX2-HaloTag in mESCs and NPCs estimated from kinetic modelling of live-cell SMT data. (**L-M**) Long residence time (τ_l_) and target-search time (τ_s_) for CBX2-HaloTag in mESCs and NPCs estimated from kinetic modelling of live-cell SMT data. (**N**) Estimated protein number per nucleus, protein number per condensate, protein concentration in nucleus, total proteins on chromatin, and stable chromatin-bound proteins for CBX2-HaloTag in ESCs and NPCs. Error represents standard deviation. Results are from three biological triplicates.

To estimate the number of Polycomb proteins per condensate, we first assessed expression levels of the fusions compared to the expression levels of endogenous proteins. Immunoblotting results confirmed that these fusion proteins are as abundant as their endogenous counterparts (**Figure S1C**). To compare the relative protein level of Polycomb proteins, we immunoblotted nuclear extracts from these cell lines. As shown in **Figure S1D,** CBX2 was undetectable while RING1B and EZH2 were most abundant. Because CBX2 was undetectable in mESCs immunoblots with HaloTag antibodies, we quantified the relative protein abundance of PRC1 and PRC2 using single-molecule HILO microscopy under the same imaging settings. Consistent with immunoblotting data, fluorescent micrographs revealed that CBX2 has the lowest protein level and RING1B and EZH2 have the highest (**Figure 1D**). Quantitative analysis showed that CBX2 was ∼40-fold less abundant than RING1B and EZH2 (**Figure 1E**). To estimate the number of proteins within condensates, we quantified RING1B by immunoblotting RING1B-HaloTag against recombinant HaloTag protein (**Figure 1F**). The RING1B concentration was estimated to be ∼203 nM per nucleus (**Figure 1G**), which is consistent with earlier reports ^48,91^. Based on the condensed fraction and number of condensates, we estimated the number of RING1B proteins per condensate and per cell (**STAR Methods**). There were ∼112×10^3^ RING1B proteins per nucleus with ∼69 RING1B proteins per condensate (**Figure 1G**). Based on the fluorescence intensity related to RING1B, we then estimated the amount of the other Polycomb fusion proteins (see **STAR Methods for details**). Strikingly, we calculated ∼2800 CBX2 proteins at a concentration of ∼5 nM within mESCs, approximating ∼3 CBX2 proteins per condensate.

The finding that ∼3 CBX2 proteins are present within each condensate prompted us to validate and further quantify this number. We performed a single-molecule photobleaching assay, where the number of photobleaching steps corresponds to the number of proteins within each condensate ^91^. We labeled CBX2-HaloTag in mESCs under two conditions: sparse labelling in which a small fraction of proteins are labelled, and full labelling where all proteins are labelled (**Figure 1H**). We performed single molecule photobleaching under single-molecule HILO microscopy. In the sparsely labeled mESCs, bleaching one spot corresponds to one photobleaching step, while in the fully labeled mESCs, completely bleaching one spot corresponds to three photobleaching steps (**Figure 1I**). Statistical analysis of multiple cells showed that there are ∼3 CBX2 proteins per condensate (**Figure 1J**). These results further support that there are ∼3 CBX2 proteins per condensate in mESCs.

According to cellular concentrations derived from the above quantification (**Figure 1G**) and our earlier *in vitro* reconstitution studies ^39^, all PRC1 proteins, except for CBX2, should not be able to form condensates on their own in cells. As we have previously shown, CBX2 forms condensates at 10 nM ^39^. We then explored whether CBX2 can form condensates at concentrations below that to recapitulate the endogenous level. An *in vitro* condensate assay carried out without crowding agents or other components revealed that CBX2 forms condensates at a concentration of 5 nM (**Figure S1E**) and that the critical concentration is close to 5 nM (**Figure S1F**). Together, these data suggested that CBX2 can form condensates at its endogenous expression level.

### Cell-type-dependent control of the number of CBX2 proteins per condensate

Because the expression of Polycomb genes is highly regulated, we asked if the number of CBX2 proteins per condensate varies in different cell types. We differentiated CBX2-HaloTag mESCs into neural progenitor cells (NPCs) (**Figure 2A**) and immunostained cells using the NPC mark of Nestin. As shown in **Figure 2B**, ∼80% of differentiated cells were Nestin-positive, and live-cell imaging confirmed that CBX2 forms condensates in NPCs (**Figure 2C**). Quantification of fluorescent signal intensity showed that CBX2 concentration in NPCs is ∼3-fold higher than in mESCs (**Figure 2D**). The condensed fraction and condensate size of CBX2 in NPCs were significantly larger than in mESCs (**Figure 2E** and **S2A**) while the number of condensates and the signal intensity ratio of condensate vs whole cell were similar (**Figure S2B-C**). We estimated that there are ∼15 proteins per condensate in NPC (**Figure 2N**), which is ∼5-fold more than in mESCs. As such, these results show that the number of CBX2 proteins per condensate is cell-type dependent.

### Cell-type-dependent regulation of CBX2 interaction with chromatin

Since there are ∼2800 molecules of CBX2 in mESCs and ∼8200 in NPCs (**Figure 2N**), we examined how many of these proteins are bound to chromatin. To this end, we performed live-cell single-molecule tracking (SMT) using single-molecule HILO microscopy (**Figure 2F**) ^24,25,37,92^. Using two different imaging modalities (**Figure 2G**), we determined the kinetic fraction and residence time of CBX2 ^24,25,37^. Under the short dark time, we measured the displacements of CBX2 in mESCs and NPCs (**Figure 2H**). The histogram of displacements was decomposed into three populations: free diffusion, confined motion, and total chromatin bound. The total chromatin-bound fraction (F_1_) of CBX2 in NPCs was larger than in mESCs (**Figure 2J**). The total number of CBX2 proteins bound to chromatin was estimated as ∼1400 in mESCs and ∼4900 in NPCs, respectively (**Figure 2N**), indicating that cellular fates dictate the number of CBX2 proteins on chromatin.

To measure the residence time and the stable chromatin-bound fraction, we performed live-cell SMT using 30-ms integration time and 170-ms dark time (**Figure 2G**). The survival probability distribution of CBX2 dwell time was fitted with a two-component exponential decay function (**Figure 2I**). The stable chromatin-bound fraction (F_s_) in NPCs was nearly 2-fold larger than that in mESCs (**Figure 2K**). There were ∼759 CBX2 proteins stably bound to chromatin in NPCs, which is ∼5-fold more than that in mESCs (**Figure 2N**). Although the number of CBX2 proteins bound to chromatin in NPCs was more than that in mESCs, their long residence time (τ_l_) was similar (**Figure 2L**). By kinetic modeling, we estimated the search time, the time for one CBX2 protein to locate its stably bound site. CBX2 took nearly 2-fold less time to locate its stable sites in NPCs than in mESCs (**Figure 2M**). Collectively, these results demonstrate that cellular states have no impact on the residence time of CBX2 but dictate its target-search time.

### Sparse CBX2 nucleates Polycomb condensates

Our quantitative analysis shows that ∼3 CBX2 proteins exist within each condensate in mESCs, which raises the question of whether sparse CBX2 can control the clustering of Polycomb proteins. To address this question, we inserted Venus-dTAG into the C-terminus of both alleles of the endogenous *Ezh2* gene in a CBX2^fl/fl^-HaloTag conditional knockout mESC line (**Figure S3A-B**). Live-cell imaging showed that EZH2 is enriched within condensates that colocalize with CBX2 condensates (**Figure 3A**). We then depleted CBX2 by administrating 4-hydroxytamoxifen (OHT) for 2 days (**Figure S3C**). The distribution of EZH2 in *Cbx2^−/−^* cells was more uniform than in *Cbx2^+/+^* cells (**Figure 3B**). Quantification analysis showed that the condensed fraction and the number of condensates of EZH2 in *Cbx2^−/−^* cells were significantly smaller than in *Cbx2^+/+^*cells (**Figure 3C-D**). To investigate whether EZH2 affects the formation of CBX2 condensates, we depleted EZH2 by administrating dTAG-13 for 24 hours (**Figure S3D**). CBX2 still formed condensates upon the depletion of EZH2 (**Figure S3E**). The condensed fraction of CBX2 was similar between wild-type and EZH2-depleted cells; however, upon the depletion of EZH2, the number of CBX2 condensates was reduced while the size of CBX2 condensates was increased (**Figure S3F**). As such, these results show that sparse CBX2 is needed for the clustering of EZH2, which in turn regulates the assembly of CBX2 condensates.

**Figure 3.**
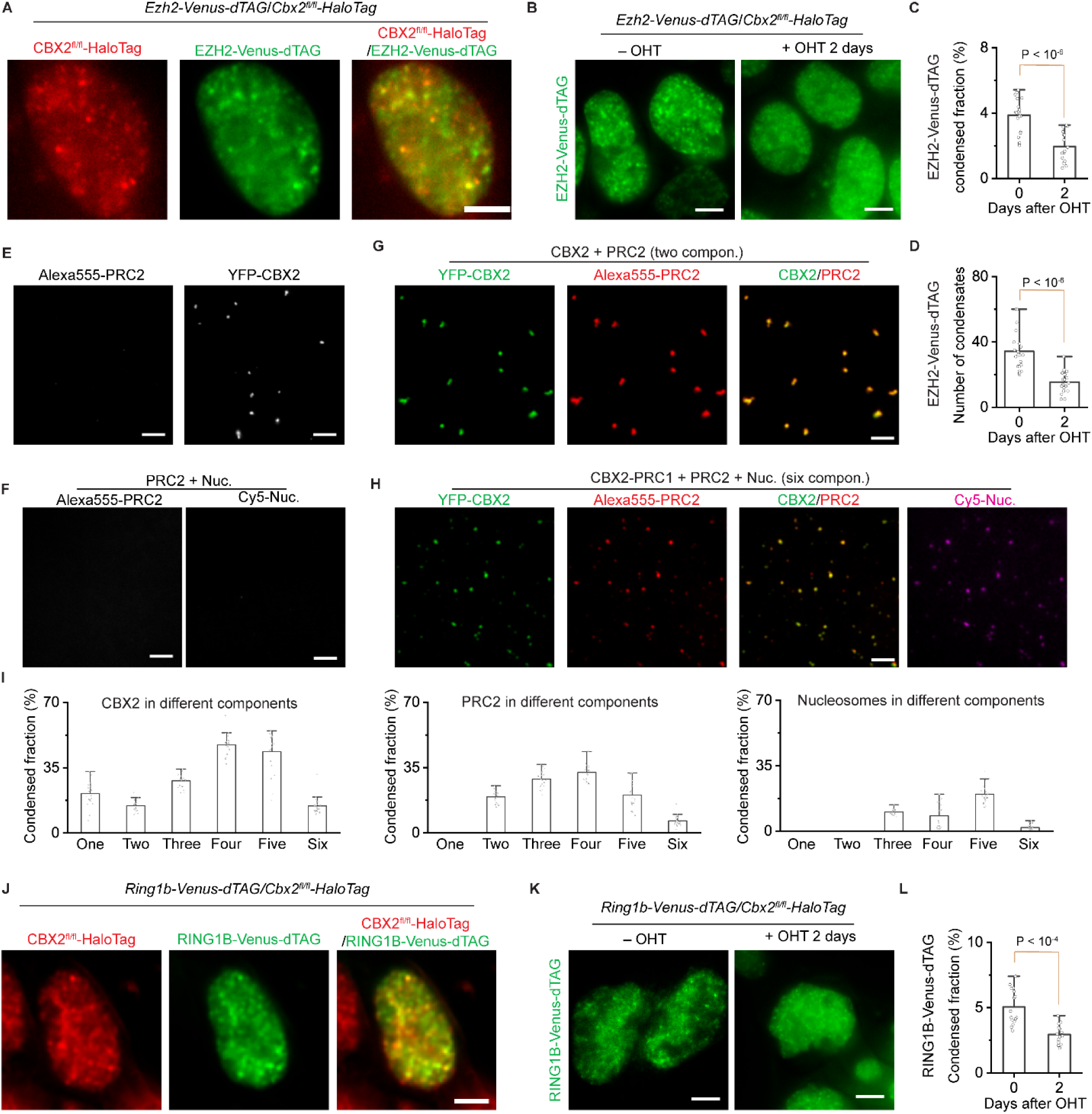
Sparse CBX2 nucleates Polycomb condensates. (**A**) Live-cell epifluorescence images of CBX2-HaloTag (red) and EZH2-Venus-dTAG (green) in *Ezh2-Venus-dTAG*/*Cbx2^fl/fl^-HaloTag* dual knockin mESC lines. Overlay image is shown. Scale bar, 5.0 µm. (**B**) Live-cell epifluorescence images of EZH2-Venus-dTAG in *Ezh2-Venus-dTAG*/*Cbx2^fl/fl^-HaloTag* dual knockin mESCs before and after depletion of CBX2. CBX2 was depleted by administrating OHT for two days. Scale bar, 5.0 µm. (**C**) Bar overlap plot of the condensed fraction of EZH2-Venus-dTAG quantified from Figure 3B. P value, student’s t-test with two-tailed distribution and two-sample unequal variance. (**D**) Bar overlap plot of the number of condensates of EZH2-Venus-dTAG quantified from Figure 3B. P value, student’s t-test with two-tailed distribution and two-sample unequal variance. (**E**) Epifluorescence images of a one-component system of YFP-CBX2 or Alexa555-labelled PRC2 consisted of EZH2, SUZ12, EED, and RbAp46/48. When defining the number of components, solvent was not counted. For the sake of simplicity, PRC2 was defined as one component. YFP-CBX2 formed condensates but Alexa555-PRC2 did not form. YFP-CBX2 was 500 nM and Alexa555-PRC2 was 50 nM. Scale bar, 5.0 µm. (**F**) Epifluorescence images of a two-component system of Alexa555-PRC2 and Cy5-nucleosomes. Alexa555-PRC2 and Cy5-nucleosomes were mixed and did not appear as condensates. Alexa555-PRC2 was 50 nM and Cy5-nucleosomes were 200 nM. Scale bar, 5.0 µm. (**G**) Epifluorescence images of condensates of a two component-system of YFP-CBX2 and Alexa555-labelled PRC2. YFP-CBX2 and Alexa555-PRC2 were mixed. PRC2 was enriched within YFP-CBX2 condensates. YFP-CBX2 was 500 nM and Alexa555-PRC2 was 50 nM. Scale bar, 5.0 µm. (**H**) Epifluorescence images of condensates of a six-component system of YFP-CBX2, Alexa555-PRC2, and Cy5-nucleosomes. The system includes CBX2 (500 nM), RING1B (500 nM), MEL18 (500 nM), PHC1 (500 nM), nucleosomes (200 nM), and PRC2 (50 nM). Individual and overlay images of CBX2 and PRC2 were shown. Images of nucleosomes were also shown. Scale bar, 5.0 µm. (**I**) Bar overlap plot of the condensed fraction of CBX2, PRC2, and nucleosomes quantified from Figure S3G. The condensed fraction was from the systems with different components. (**J**) Live-cell epifluorescence images of CBX2-HaloTag (red) and RING1B-Venus-dTAG (green) in *Ring1b-Venus-dTAG*/*Cbx2^fl/fl^-HaloTag* dual knockin mESC lines. Overlay images were shown. Scale bar, 5.0 µm. (**K**) Live-cell epifluorescence images of RING1B-Venus-dTAG in *Ring1b-Venus-dTAG/Cbx2^fl/fl^-HaloTag* dual knockin mESCs upon the depletion of CBX2. CBX2 was depleted by administrating OHT for two days. Scale bar, 5.0 µm. (**L**) Bar overlap plot of the condensed fraction of RING1B-Venus-dTAG quantified from Figure 3K. P value, student’s t-test with two-tailed distribution and two-sample unequal variance.

To investigate the molecular mechanism underlying the clustering of PRC2 by CBX2, we performed *in vitro* condensate formation assays. We labeled the recombinant PRC2 complex, which includes full-length EZH2, SUZ12, EED, and RbAp46/48, using Alexa555 fluorescence dyes. We found that the labeled PRC2 alone did not form condensates while CBX2 formed condensates independently (**Figure 3E**). Since nucleosomes are the substrate of PRC2, we asked whether nucleosomes could help the formation of PRC2 condensates. In the heterogenous system, PRC2 and nucleosomes still did not form condensates (**Figure 3F**). We then mixed CBX2 and PRC2 (thereafter, for the sake of simplicity, the PRC2 complex was referred to as one component) and found PRC2 was partitioned into CBX2 condensates (**Figure 3G**). Since the assembly of CBX2-PRC1 condensates is composition-dependent ^39^, we investigated whether increasing the number of components within the system regulates the enrichment of PRC2 into condensates. Adding nucleosomes to the two-component system increased the enrichment of PRC2 into condensates and adding RING1B (the four-component system) further increased the enrichment (**Figure 3H-I** and **S3G-H**). Adding MEL18 (the five-component system) and PHC1 (the six-component system) decreased the enrichment of PRC2 into condensates. Thus, these results suggest that PRC2 is also partitioned into phase-separated CBX2 condensates in a composition-dependent manner.

Recent studies have shown that CBX2 drives the formation of CBX-PRC1 condensates ^36,38,39,47^. The depletion of CBX2 resulted in a dispersion of condensates of CBX7 and PHC1 in mESCs ^39^, the components of the CBX-PRC1 complexes. Since RING1B is the core component of both CBX-PRC1 and variant PRC1, we investigated whether sparse CBX2 also regulates the clustering of RING1B. We inserted Venus-dTAG into the C-terminus of one allele of the endogenous *Ring1b* gene in CBX2^fl/fl^-HaloTag mESC lines (**Figure S3A-B**). Live-cell imaging showed that RING1B forms condensates that colocalize with CBX2 condensates (**Figure 3J**). The depletion of CBX2 resulted in a significant reduction in the condensed fraction of RING1B (**Figure 3K-L**). This data indicates that sparse CBX2 also controls the assembly of PRC1 condensates.

### Sparse CBX2 demarcates the spatial boundaries of facultative heterochromatin and controls the H3K27me3 abundance

Facultative heterochromatin is characterized by high levels of Polycomb proteins and H3K27me3 ^1–3^. As sparse CBX2 nucleates Polycomb proteins, we reasoned that CBX2 could demarcate the spatial boundaries of facultative heterochromatin. To this end, we depleted CBX2 in *EZH2-Venus-dTAG/CBX2^fl/fl^-HaloTag* cells by administrating OHT (**Figure 4A**). Following fixation, DNA was stained by Hoechst. Confocal imaging showed that EZH2 in *Cbx2^+/+^* cells is excluded from DNA-dense regions that are constitutive heterochromatin; however, upon the depletion of CBX2, EZH2 migrated onto DNA-dense regions. We quantified the ratio between the fluorescence intensity of EZH2 and the Hoechst signal of DNA-dense regions. EZH2 was significantly more enriched in DNA-dense regions in *Cbx2^−/−^* cells than in *Cbx2^+/+^* cells (**Figure 4B**). We then studied whether sparse CBX2 is needed for the spatial distribution of PRC1 proteins. Like EZH2, the depletion of CBX2 similarly resulted in a migration of CBX7, PHC1, and RING1B to DNA-dense regions (**Figure S4A-B**). Thus, these results show that sparse CBX2 is needed for the control of the spatial boundaries of Polycomb proteins.

**Figure 4.**
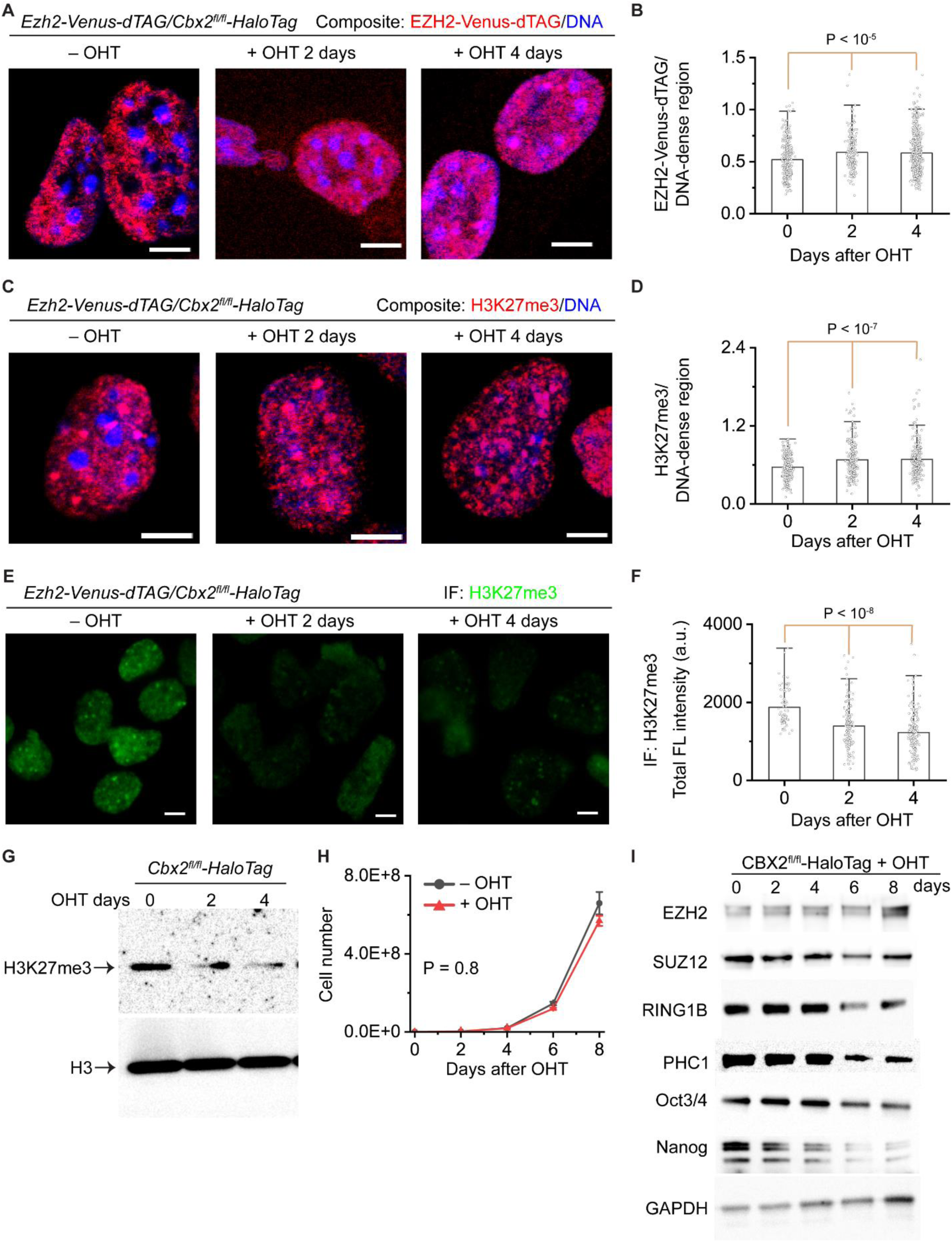
Sparse CBX2 demarcates the spatial boundaries of facultative heterochromatin and controls the H3K27me3 level. (**A**) Representative confocal images of EZH2-Venus-dTAG in *Ezh2-Venus-dTAG/Cbx2^fl/fl^-HaloTag* mESCs. *Cbx2* is depleted by administrating OHT. DNA was stained by Hoechst. Overlay images are shown. EZH2-Venus-dTAG is red and DNA is blue. Scale bar, 5.0 µm. (**B**) Bar overlap plot of the fluorescence intensity ratio of EZH2-Venus-dTAG to DNA dense regions quantified from Figure 4A. P value, student’s t-test with two-tailed distribution and two-sample unequal variance. (**C**) Representative confocal images of H3K27me3 immunostained by anti-H3K27me3 antibodies in *Ezh2-Venus-dTAG*/*Cbx2^fl/fl^-HaloTag* mESCs. *Cbx2* is depleted by administrating OHT. DNA was stained by Hoechst. Overlay images are shown. H3K27me3 is red and DNA is blue. Scale bar, 5.0 µm. (**D**) Bar overlap plot of the fluorescence intensity ratio of H3K27me3 to DNA dense regions quantified from Figure 4C. P value, student’s t-test with two-tailed distribution and two-sample unequal variance. (**E**) Representative epifluorescence images of H3K27me3 in *Ezh2-Venus-dTAG/Cbx2^fl/fl^-HaloTag* mESCs treated with or without OHT. H3K27me3 was immunostained by anti-H3K27me3 antibodies. Images were taken by using the same settings. Scale bar, 5.0 µm. (**F**) Bar box plot of the fluorescence intensity of H3K27me3 quantified from Figure 4E. P value, student’s t-test with two-tailed distribution and two-sample unequal variance. (**G**) Immunoblotting of histones extracted from *Ezh2-Venus-dTAG/Cbx2^fl/fl^-HaloTag* mESCs treated with or without OHT. Blots were probed by anti-H3K27me3 antibodies and anti-H3 antibodies. (**H**) Growth rate of *Ezh2-Venus-dTAG/Cbx2^fl/fl^-HaloTag* mESCs treated with or without OHT. P value, student’s t-test with two-tailed distribution and two-sample unequal variance. (**I**) Immunoblotting of PRC2 and PRC1 proteins as well as the pluripotency factors Oct3/4 and Nanog extracted from *Ezh2-Venus-dTAG/Cbx2^fl/fl^-HaloTag* mESCs treated with or without OHT. GAPDH was used for loading control.

Since CBX2 regulates the spatial boundaries of PRC2, we then speculated that CBX2 is also needed for the control of the spatial distribution of H3K27me3. We immunostained H3K27me3 in *Cbx2^+/+^*and *Cbx2^−/−^* cells. Confocal imaging showed that H3K27me3 in *Cbx2^+/+^* cells is excluded from DNA-dense regions; however, in *Cbx2^−/−^* cells, H3K27me3 is distributed into DNA-dense regions (**Figure 4C**). Quantitative analysis showed that the ratio of H3K27me3 to DNA-dense regions in *Cbx2^−/−^* cells is significantly larger than that in *Cbx2^+/+^* cells (**Figure 4D**). Overall, these data show that CBX2 is needed for the spatial distribution of facultative heterochromatin.

Given that CBX2 controls the spatial distribution of Polycomb proteins and H3K27me3, we hypothesized that CBX2 could globally regulate the level of H3K27me3. To test this, we performed quantitative epifluorescence imaging of H3K27me3 in *Cbx2^+/+^* and *Cbx2^−/−^* cells (**Figure 4E-F**). Quantitative analysis showed that the H3K27me3 level in *Cbx2^−/−^* cells are ∼70% of that in *Cbx2^+/+^* cells. We then performed immunoblotting of histone extracts from *Cbx2^+/+^*and *Cbx2^−/−^* cells and found that the level of H3K27me3 in *Cbx2^−/−^* cells are less than that in *Cbx2^+/+^*cells (**Figure 4G**). These results show that CBX2 regulates the level of H3K27me3.

The above observations could be resulted from altered cell proliferation or pluripotency. To discern this, we performed cellular proliferation assays and found that the growth rate of *Cbx2^−/−^* cells shows no significant difference compared to that of *Cbx2^+/+^* cells (**Figure 4H**). We then investigated whether CBX2 influences the protein level of PRC1 and PRC2. Immunoblotting showed that the protein level of both PRC1 and PRC2 is similar for the first 4-days of culture; however, except for EZH2, long-term culture of *Cbx2^−/−^* cells results in a reduction in their protein level (**Figure 4I**). The levels of the markers of pluripotency proteins, Oct3/4 and Nanog, were similar for the first 4-day culture but was reduced beyond 4 days (**Figure 4I**). Our results suggest that CBX2 is not needed to maintain the levels of Polycomb proteins and pluripotency in short-term culture but is needed for their long-term maintenance.

### CBX2 loss results in genome-wide redistribution of H3K27me3 that is predicted by genomic localization of CBX2

To determine where in the genome H3K27me3 is lost upon CBX2 knockout, we performed H3K27me3 CUT&Tag in duplicates over the time course of OHT treatment (0, 2, 4, and 6 days). Upon loss of CBX2, we see a decrease in H3K27me3 enrichment at known H3K27me3 domains (**Figure S5A-B**). To systematically determine changes in H3K27me3 domains genome-wide, we defined H3K27me3 domains for each time point, combined all the domain definitions, and performed the “Disjoin” operation^93^ to identify non-overlapping segments that would encompass H3K27me3 domains from all time points. We then performed k-means (k=6) clustering of the H3K27me3 changes over the time points for the non-overlapping segments. Clusters 1 to 6 are arranged in descending order of the decrease in H3K27me3 due to the loss of CBX2 (**Figure 5A-B, S5C**). Clusters 1-3 showed a decrease in H3K27me3, H3K27me3 levels did not change much in Cluster 4, whereas Clusters 5 and 6 showed a slight gain in H3K27me3 levels (**Figure S5C**). The segments that lost H3K27me3 in CBX2 KO had high overlap with WT H3K27me3 domains, whereas Clusters 5 and 6 show a slight gain in H3K27me3 had low overlap with WT domains (**Figure S5D**). Similar to the imaging experiments, these results suggest a shift in H3K27me3 enrichment from known regions to new regions due to the loss of CBX2. Segments comprising cluster 1, which had the largest loss in H3K27me3 due to CBX2 loss, had significant overlap with PRC2 nucleation sites defined in ref.^19^ (**Figure S5E**). Furthermore, the overlap with CpG islands and bivalent promoters was highest for Cluster 1 and decreased monotonically from Cluster 1 to Cluster 6 (**Figure S5F**). Thus, CBX2 loss results in a decrease in H3K27me3 at sites of PRC2 nucleation. We next asked if the sites that lose H3K27me3 upon CBX2 knockout are also enriched for CBX2 binding. We performed CBX2 CUT&RUN and plotted CBX2 enrichment at clusters defined based on H3K27me3 change upon CBX2 knockout. Notably, we observe CBX2 enrichment to be highest at Cluster 1, which has the biggest loss in H3K27me3 upon loss of CBX2, with CBX2 enrichment going down from Cluster 1 to Cluster 6 (**Figure 5C-D**). To ask whether CBX2 loss affects EZH2 occupancy, we performed CUT&RUN of EZH2 in wild-type and CBX2 knockout cells. EZH2 decreased from Cluster I to Cluster 6, and upon CBX2 loss, was reduced the most for Clusters 1-3 (**Figure 5E-F**). Thus, the change in H3K27me3 and EZH2 upon loss of CBX2 correlates with the enrichment of CBX2 in WT cells. We can conclude that CBX2 binds at PRC2 nucleation sites and may help with PRC2 nucleation to maintain proper H3K27me3 distribution in the genome.

**Figure 5.**
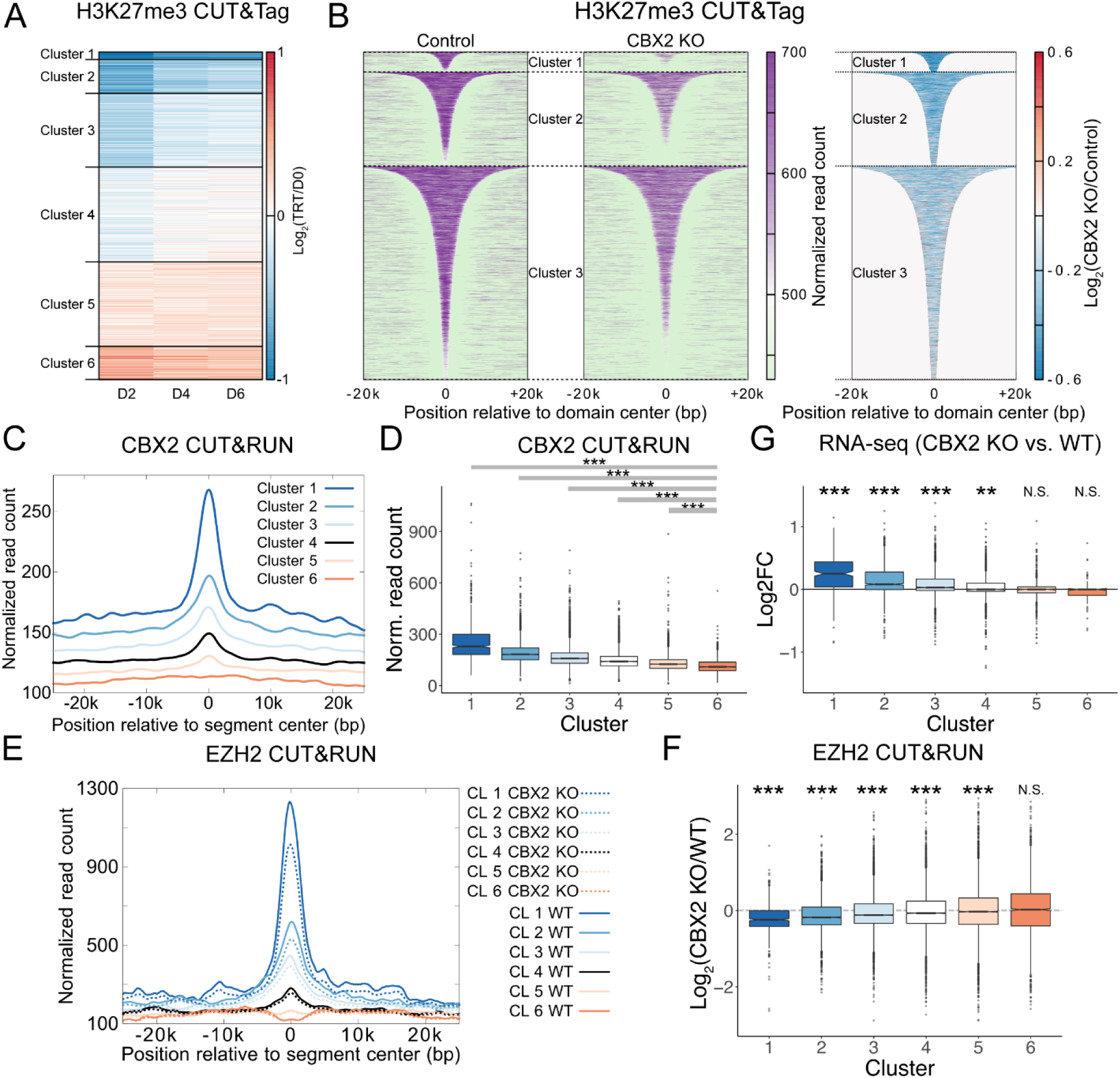
Effect of CBX2 loss on H3K27me3 domains. (**A**) Changes in H3K27me3 CUT&Tag enrichment at unique segments defined by Disjoin. Log2 enrichment over Day 0 is plotted as a heatmap. (**B**) H3K27me3 enrichment at Day 0 (Control, left) and Day 2 of OHT treatment (CBX2 KO, middle) are plotted for domains from the clusters defined in (A). Log2 ratio of H3K27me3 enrichment in CBX2 KO over Control within the H3K27me3 domains is plotted as a heatmap (right). (**C**) Normalized read counts from CBX2 CUT&RUN averaged over segments of each of the six clusters defined in (A) are plotted relative to the center of the segments. (**D**) CBX2 CUT&RUN enrichment was calculated at each H3K27me3 segment defined in (A) and the distribution for each cluster is shown as a boxplot. Wilcox rank sum test was used to calculate the p-value for the comparison between each cluster and cluster 6 and p-values were adjusted for multiple comparisons using the Bonferroni correction. *** denotes p < 2.2e-16. (**E**) Normalized read counts from EZH2 CUT&RUN averaged over segments of each of the six clusters defined in (A) are plotted relative to the center of the segments. (**F**) Log2 ratio of EZH2 CUT&RUN enrichment in WT to that in CBX2 KO was calculated at each H3K27me3 segment defined in (A), and the distribution for each cluster is shown as a boxplot. Wilcox rank sum test was used to calculate the p-value with the null hypothesis that the log2 fold change for each cluster is 0, and p-values were adjusted for multiple comparisons using the Bonferroni correction. *** denotes p < 2.2e-16, N.S. denotes not significant (p>0.05). (**G**) The distribution of log2 fold change in TPM values for genes overlapping segments plotted in (A) are shown as boxplots. Wilcox rank sum test was used to calculate the p-value with the null hypothesis that the log2 fold change for each cluster is 0, and p-values were adjusted for multiple comparisons using the Bonferroni correction. *** denotes p < 2.2e-16, ** denotes p= 1.9e-08, N.S. denotes not significant (p>0.05).

We next asked if there is a loss of repression that correlates with the loss of H3K27me3. We performed RNA-seq on day 0 and day 2 after OHT treatment and calculated log2 fold-change for genes that overlap the 6 clusters defined based on changes in H3K27me3 upon loss of CBX2. We observe an upregulation of genes whose transcription start sites (TSSs) lie within regions in clusters 1, 2, and 3, but not in clusters 4, 5, and 6 (**Figure 5G**). In conclusion, genes that lose H3K27me3 due to the loss of CBX2 also feature an increase in expression, pointing to a loss of silencing due to the loss of CBX2. Taking all the genomic analysis together, we can conclude that CBX2 binds at PRC2 nucleation sites, helps maintenance of H3K27me3 and repression of transcription. Loss of CBX2 leads to relocation of H3K27me3 from its regular targets in mESC, presumably due to weakened nucleation of PRC2.

### CBX2 is needed for proper cellular differentiation

Despite the low abundance of CBX2 in mESCs, CBX2 regulates H3K27me3 deposition and spatial distribution and represses developmental regulators, suggesting CBX2 could control cell-fate transition. We then investigated whether *Cbx2^−/−^* mESCs are defective in cellular differentiation by forming embryoid bodies (EBs) in the presence or absence of retinoic acid. *Cbx2^+/+^* and *Cbx2^−/−^* cells could form EBs, but the EB size of *Cbx2^−/−^* cells was smaller than that of *Cbx2^+/+^*cells (**Figure S6A-B**). We then plated EBs on gelatin-coated plates to monitor outgrowth, which is indicative of robust differentiation. After 24 hours of culture, there was almost no outgrowth from EBs of *Cbx2^−/−^* compared to *Cbx2^+/+^* (**Figure 6A** and **S6C**). To study whether the depletion of CBX2 affects the development of lineage-specific cell types, we differentiated both *Cbx2^+/+^* and *Cbx2^−/−^* cells into NPCs. Both the total number of cells and the number of Nestin-positive cells derived from *Cbx2^−/−^* cells were significantly lower than that derived from *Cbx2^+/+^* cells (**Figure 6B-C** and **S6D**). Thus, these results show that CBX2 is needed for the proper differentiation of mESCs.

**Figure 6.**
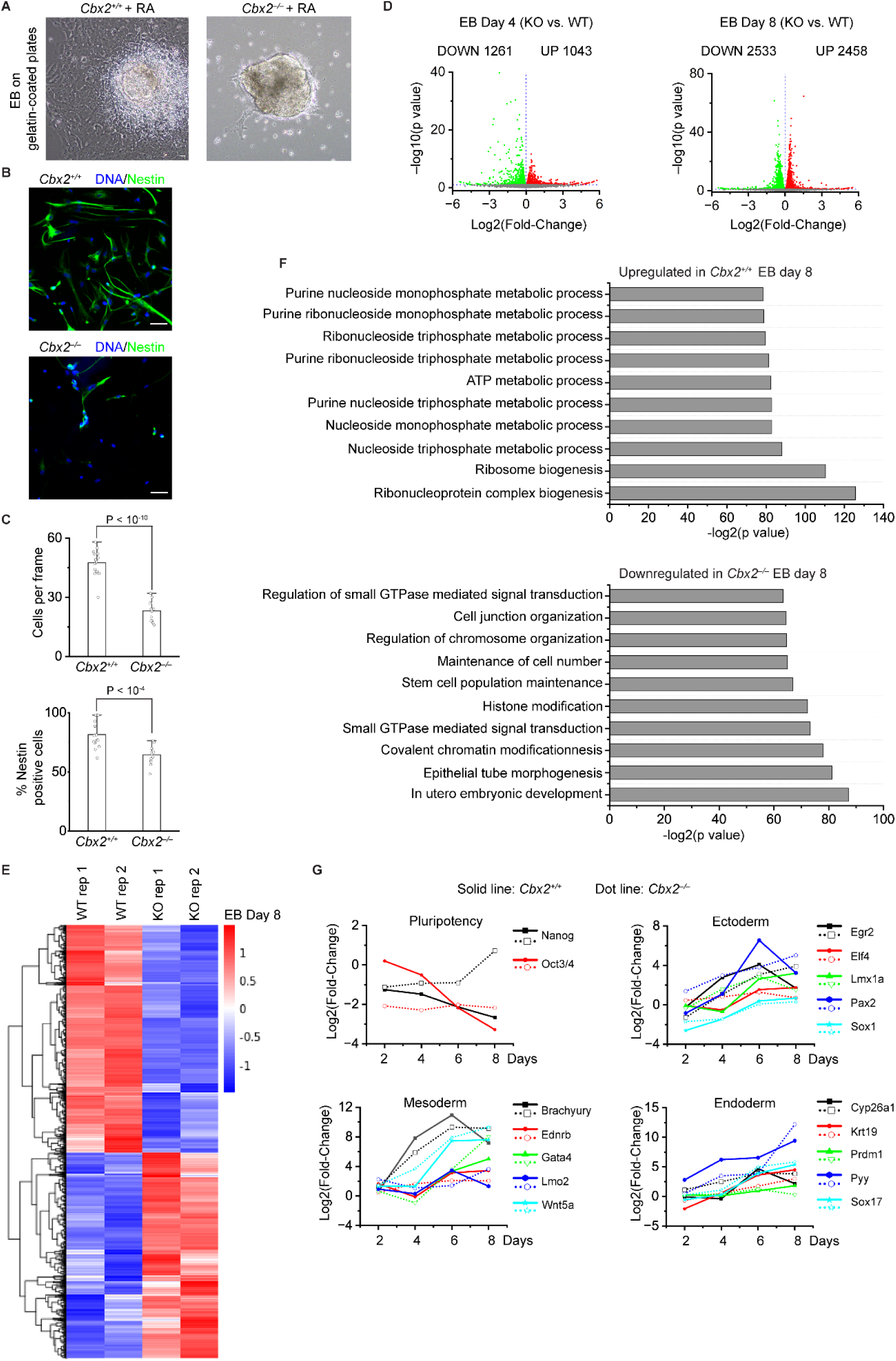
CBX2 is needed for proper cellular differentiation. (**A**) Phase-contrast images of the outgrowth of day-8 EBs on gelatin-coated plates for 24 hours for wild-type (*Cbx2^+^/^+^*) and knockout (*Cbx2^−/−^*) cells in the presence of retinoic acid (RA). Scale bar, 100 µm. (**B**) Representative epifluorescence images of immunostained NPCs derived from wild-type (*Cbx2^+^/^+^*) and knockout (*Cbx2^−/−^*) cells under the same differentiation conditions. Cells were immunostained by anti-Nestin antibodies. DNA was stained by Hoechst. Overlay images are shown. Nestin, green color. DNA, blue color. Scale bar, 50 µm. (**C**) Number of cells per frame and percentage of Nestin-positive cells. Cells were derived from wild-type (*Cbx2^+^/^+^*) and knockout (*Cbx2^−/−^*) cells under the same differentiation conditions. P value, student’s t-test with two-tailed distribution and two-sample unequal variance. (**D**) Volcano plot of differentially expressed genes (DEGs) between wild-type (*Cbx2^+^/^+^*) and knockout (*Cbx2^−/−^*) from day-4 (left) and day-8 (right) EBs, with numbers of significantly up-regulated (red) and down-regulated (green) genes (red dots, p<0. 1, |log2 fold change|>1) shown. (**E**) The hierarchical clustering analysis of the DEGs between wild-type (*Cbx2^+^/^+^*) and knockout (*Cbx2^−/−^*) from day-8 EBs. RNA expression values (RPKM, reads per kilobase per million reads) are represented by z-score across samples. (**F**) Gene ontology (GO) analysis of the up-regulated and down-regulated genes from day-8 EBs with the top 10 significant GO terms and p value displayed. (G) Transcriptional activities of pluripotent genes and lineage-specific genes for differentiation of wild-type (*Cbx2^+^/^+^*) and knockout (*Cbx2^−/−^*) cells for different days.

To investigate whether the defective differentiation of *Cbx2^−/−^* cells is caused by dysfunctional transcriptional programs, we analyzed transcriptomics of day-4 and day-8 EBs from *Cbx2^+/+^* and *Cbx2^−/−^* cells (**Figure 6D-E** and **S6E**). ∼1000 genes in day-4 EBs were up-regulated and ∼1200 genes were down-regulated when comparing *Cbx2^+/+^* and *Cbx2^−/−^* cells. Culturing of EBs to day-8 led to ∼5000 genes that are either up- or down-regulated between the WT and KO groups. Gene ontology (GO) analysis showed that the down-regulated genes are enriched for embryonic development, morphogenesis, stem cell maintenance, and histone modification, while the up-regulated genes relate to metabolic processes (**Figure 6F**). We further analyzed pluripotent genes and lineage-specific markers by RT-qPCR. Both pluripotent genes and lineage-specific markers are mis-regulated during cellular differentiation of *Cbx2^−/−^* cells (**Figure 6G**). As such, our results show that CBX2 is needed for maintaining proper transcriptional programs during cellular differentiation.

### LLPS of CBX2 regulates the spatial distribution of H3K27me3

Since CBX2 regulates the spatial distribution of H3K27me3, we asked whether the LLPS of CBX2 contributes to this process. To address this, we generated a loss-of-function variant, CBX2^DM^, where the positively charged amino acid residues within the ATL and HPCR motifs are replaced with alanine residues (**Figure 7A**). To investigate whether CBX2^DM^ forms condensates, we purified the recombinant CBX2^DM^ and performed *in vitro* condensate formation assays. CBX2^DM^ only formed condensates at a critical concentration that is over 100-fold higher than CBX2 (**Figure 7B** and **S7B**). By using CRISPR/Cas9, we mutated the same set of amino acid residues to generate *Cbx2^DM^-HaloTag (HT)* mESC lines (**Figure 7C** and **S7C-D**). CBX2^DM^-HaloTag did not form condensates in cells (**Figure 7D**). To interrogate whether the LLPS ability of CBX2 is needed for demarcating the spatial boundaries of H3K27me3, we immunostained *Cbx2^DM^-HaloTag* and *Cbx2^+/+^* cells by using H3K27me3 antibodies and Hoechst. Confocal micrographs showed that H3K27me3 has higher enrichment on DNA-dense regions in *Cbx2^DM^-HaloTag* cells compared to *Cbx2^+/+^*cells (**Figure 7E**), similar to *Cbx2^−/−^* cells. Quantitative analysis showed that the intensity ratio of H3K27me3 to DNA-dense regions in *Cbx2^DM^-HaloTag* cells is significantly larger than that in *Cbx2^+/+^* cells (**Figure 7F**). Thus, these results show the LLPS of CBX2 is needed for the spatial distribution of H3K27me3.

**Figure 7.**
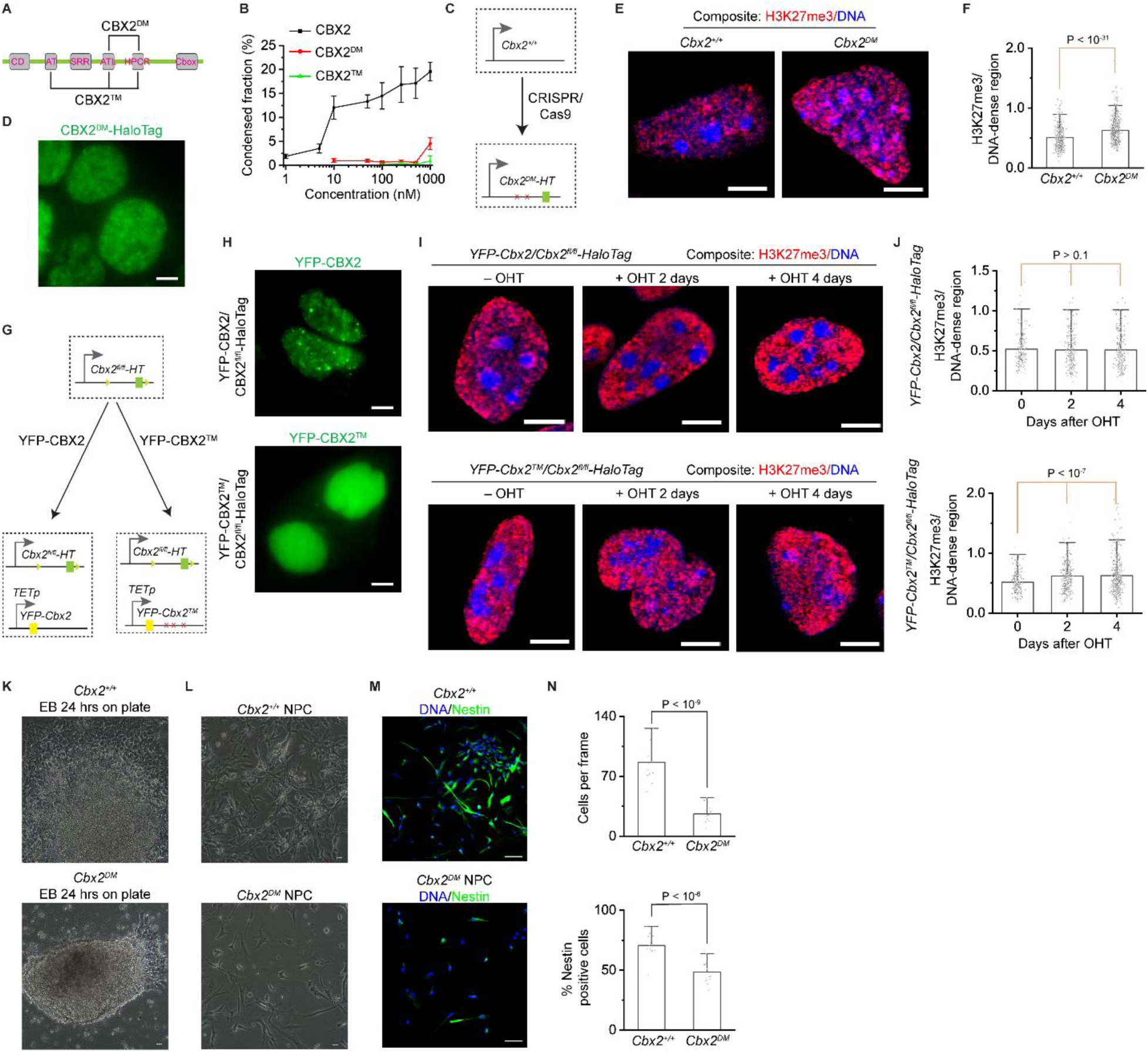
LLPS of CBX2 demarcates the spatial boundaries of facultative heterochromatin and regulates cellular differentiation. (**A**) Schematic representation of the secondary structure of CBX2. CBX2^DM^ is a variant of CBX2 where the Arg and Lys residues within ATL and HPCR have been substituted with Ala. CBX2^TM^ is a variant of CBX2 where the Arg and Lys residues within AT, ATL, and HPCR have been substituted with Ala. CD, chromodomain. AT, AT-hook motif. ATL, AT-hook-like motif. HPCR, highly positively charged region. Cbox, chromobox. (**B**) Plot of the condensed fraction of CBX2, CBX2^DM^, and CBX2^TM^ versus different protein concentrations. (**C**) Schematic representation of the generating of *Cbx2^DM^-HaloTag* (for the sake of simplicity, *Cbx2^DM^-HaloTag* sometimes was referred to as *Cbx2^DM^*) mESC lines by CRISPR/Cas9. (**D**) Live-cell epifluorescence images of CBX2^DM^-HaloTag in *Cbx2^DM^-HaloTag* mESCs. Scale bar, 5.0 µm. (**E**) Representative confocal images of H3K27me3 and DNA in wild-type and *Cbx2^DM^-HaloTag* mESCs. H3K27me3 was stained by anti-H3K27me3 antibodies, and DNA was stained by Hoechst. Composite images are shown. H3K27me3 is red, and DNA is blue. Scale bar, 5.0 µm. (**F**) Box overlap plot of the fluorescence intensity ratio of H3K27me3 to DNA dense regions quantified from Figure 7E. P value, student’s t-test with two-tailed distribution and two-sample unequal variance. (**G**) Schematic representation of the functional complementation assay. *YFP-Cbx2* and *YFP-Cbx2^TM^*were stably integrated into the genome of *Cbx2^fl/fi^-HaloTag* mESCs, respectively. (**H**) Live-cell epifluorescence images of YFP-CBX2 in *YFP-Cbx2*/*Cbx2^fl/fi^-HaloTag* mESCs and YFP-CBX2^TM^ in *YFP-Cbx2^TM^*/*Cbx2^fl/fi^-HaloTag* mESCs, respectively. Scale bar, 5.0 µm. (**I**) Representative confocal images of H3K27me3 and DNA in *YFP-Cbx2/Cbx2^fl/fl^-HaloTag* and *YFP-Cbx2^TM^/Cbx2^fl/fl^-HaloTag* mESCs, respectively. Cells were treated with or without OHT for different time periods. Composite images are shown with DNA being blue color and H3K27me3 as red color. H3K27me3 was immunostained by anti-H3K27me3 antibodies. DNA was stained by Hoechst. Scale bar, 5.0 µm. (**J**) Bar overlap plot of the fluorescent intensity ratio of H3K27me3 to DNA-dense regions in *YFP-Cbx2/Cbx2^fl/fl^-HaloTag* and *YFP-Cbx2^TM^/Cbx2^fl/fl^-HaloTag* mESCs quantified from Figure 7I. (**K**) Phase-contrast images of the outgrowth of day-8 EBs derived from wild-type and *Cbx2^DM^-HaloTag* cells on gelatin-coated plates for 24 hours. Scale bar, 100 µm. (**L**) Phase-contrast images of NPC derived from wild-type and *Cbx2^DM^-HaloTag* cells. Scale bar, 100 µm. (**M**) Representative epifluorescence images of immunostained NPCs derived from wild-type and *Cbx2^DM^-HaloTag* cells. Cells were immunostained by anti-Nestin antibodies. DNA was stained by Hoechst. Overlay images are shown with Nestin being green and DNA being blue. Scale bar, 100 µm. (**K**) Bar overlap plot of the number of cells per frame and percentage of Nestin-positive cells. P value, student’s t-test with two-tailed distribution and two-sample unequal variance.

To further study whether the LLPS of CBX2 is needed for the spatial distribution of H3K27me3, we performed functional complementation assays. We generated another loss-of-function variant, CBX2^TM^, where the positively charged amino acid residues within AT, ATL, and HPCR are replaced with alanine residues (**Figure 7A**). CBX2^TM^ also formed condensates at a critical concentration over 100-fold higher than CBX2 (**Figure 7B** and **S7A-B**). We then stably integrated *YFP-Cbx2* and *YFP-Cbx2^TM^* into *Cbx2^fl/fl^-HT* (HaloTag) mESCs (**Figure 7G**). The expression level of the transgenes was under the control of a TET-response promoter. Since the protein level of endogenous CBX2 is extremely low, these fusion genes were expressed under basal levels without doxycycline induction. Live-cell imaging showed that YFP-CBX2^TM^ does not form condensates while YFP-CBX2 forms condensates as expected (**Figure 7H**), indicating that CBX2^TM^ does not phase separate in cells.

To investigate whether the LLPS of CBX2 affects the spatial distribution and the level of H3K27me3, we depleted endogenous CBX2 in *YFP-Cbx2^TM^*/*Cbx2^fl/fl^-HaloTag* and *YFP-Cbx2*/*Cbx2^fl/fl^-HaloTag* cells by administering OHT. These cells were immunostained by H3K27me3 and Hoechst. Confocal micrographs showed that upon the depletion of CBX2, H3K27me3 is excluded from DNA-dense regions in *Cbx2^fl/fl^-HaloTag* cells complemented with *YFP-Cbx2* but migrates onto DNA-dense regions in *Cbx2^fl/fl^-HaloTag* cells complemented with *YFP-Cbx2^TM^* which is defective in phase separation (**Figure 7I**). Quantitative analysis showed that the intensity ratio of H3K27me3 to DNA-dense regions was similar between *YFP-Cbx2*/*Cbx2^fl/fl^-HaloTag* cells treated with or without OHT (**Figure 7J**). However, the intensity ratio of H3K27me3 on DNA-dense regions in *YFP-Cbx2^TM^*/*Cbx2^fl/fl^-HaloTag* cells treated with OHT was significantly larger than that without adding OHT, suggesting that CBX2^TM^ is defective in demarcating the spatial distribution of H3K27me3. To investigate whether the LLPS of CBX2 is required for maintaining H3K27me3 levels, we quantified the H3K27me3 level in *YFP-Cbx2^TM^*/*Cbx2^fl/fl^-HaloTag* and *YFP-Cbx2*/*Cbx2^fl/fl^-HaloTag* cells by administering OHT. YFP-CBX2 could restore the level of H3K27me3 but YFP-CBX2^TM^ could not (**Figure S7E-F**). Thus, these results further support that the LLPS of CBX2 is needed for the spatial distribution and the level of H3K27me3.

### LLPS of CBX2 regulates cellular differentiation

To investigate whether the LLPS of CBX2 is needed for cellular differentiation, we differentiated *Cbx2^DM^*and *Cbx2^+/+^* cells into EBs and then plated EBs on plates. 24 hours after plating, the outgrowth of *Cbx2^DM^* EBs was nearly undetectable, which is similar to *Cbx2^−/−^* EBs. To interrogate whether the LLPS of CBX2 is needed for differentiating mESCs into specific lineages, we differentiated *Cbx2^DM^* and *Cbx2^+/+^*cells into NPCs. *Cbx2^DM^* cells were defective in the NPC generation (**Figure 7L-M**). Quantitative analysis showed that the number of cells derived from *Cbx2^DM^*cells was significantly lower than that derived from *Cbx2^+/+^* cells (**Figure 7N**). The number of Nestin-positive cells derived from *Cbx2^DM^* cells was also significantly less than that derived from *Cbx2^+/+^* cells (**Figure 7N**). As such, these data show that the LLPS of CBX2 is needed for proper cellular differentiation.

## DISCUSSION

Our work provides a previously unappreciated biological framework: low-abundant proteins can execute functionalities by nucleating many other proteins to form non-stoichiometric condensates by phase separation. Here, we leverage Polycomb proteins as a paradigm to uncover this framework. Using live-cell single-molecule imaging, genetic engineering, and molecular and genomic approaches, we found that sparse CBX2 proteins nucleate many proteins of PRC1 and PRC2 to form non-stoichiometric silencing condensates, which in turn promote the facultative heterochromatinization of Polycomb target genes by facilitating H3K27me3 deposition and demarcating the spatial distribution of facultative heterochromatin. The perturbation of CBX2 phase separation results in the defective formation of facultative heterochromatin and impaired cell differentiation. Our findings yield a phase-separation-based mechanism of facultative heterochromatinization of Polycomb target genes. Furthermore, the discovery that proteins with low abundance nucleate supramolecular complexes to form phase-separated functional condensates can be generalized to other low-abundant proteins in biology.

### Sparse CBX2 nucleates Polycomb proteins to form silencing condensates

The composition-dependent control of condensate assembly by phase separation is an emerging concept underlying the biophysical properties of Polycomb condensates ^39,47^. CBX2 is an intrinsically disordered protein with alternating positively and negatively charged regions that drive the formation of homo-typic and hetero-typic CBX-PRC1 condensates ^36,38,39,47^. We show that PRC2, which does not form condensates *in vitro*, is partitioned into CBX2-PRC1 condensates in a composition-dependent manner, suggesting that CBX2-PRC1 controls the compartmentalization of PRC2. Earlier studies have shown that H3K27me3, the PRC2 catalytic product, recruits CBX-PRC1 complexes to chromatin through CBX subunits recognizing H3K27me3 ^92,94,95^. Together, these results suggest a new model by which PRC2 and CBX-PRC1 coordinate to form a feedback loop through compartmentalizing PRC2 by CBX2-PRC1 and recognizing H3K27me3 by CBX-PRC1.

In mESCs, CBX2 is at a low substoichiometric level compared to other Polycomb proteins within condensates. However, despite its low level, CBX2 is essential for nucleating PRC1 and PRC2 to form silencing condensates, which demarcate the spatial boundaries of facultative heterochromatin and facilitate H3K27me3 deposition. Our results suggest the following model. One CBX2 protein is stably bound to chromatin, which nucleates the binding of two more CBX2 proteins through homo-typic interactions. Three CBX2 proteins then recruit CBX-PRC1 and PRC2 through heterotypic interactions to form Polycomb condensates. Together, our results imply that sparse CBX2 acts as a seed to nucleate CBX-PRC1 and PRC2 to form silencing condensates, which are critical for the facultative heterochromatinization of Polycomb target genes.

We also show that Polycomb condensates are reprogrammed during cell differentiation. The number of condensates remains similar, but the number of CBX2 proteins per condensate increases from ∼3 in mESCs to ∼15 in NPCs, and the number of stably bound CBX2 proteins per cell increases from ∼135 in mESCs to ∼759 in NPCs. The change in the number of CBX2 per condensate confers contextually specific biophysical properties for condensates, which have broad implications for controlling cell-type specific transcription programs. For instance, the silencing of Polycomb targets in pluripotent cells should be more plastic in order to rapidly respond to differential signals whereas the differentiated cellular states require relatively stable silencing. Recent studies have shown that the exact composition of Polycomb condensates affects the stability of condensates ^39,47^. Since Polycomb proteins are developmentally regulated ^10^, we propose that the exact composition within condensates during development will dictate the specific material properties of condensates, which are needed for specific cell-type-appropriate transcription programs.

### A phase-separation-based model underlying facultative heterochromatinization of Polycomb target genes

Our results uncover a new functional role of sparse CBX2 in the facultative heterochromatinization of Polycomb target genes, which is achieved through nucleating many proteins of PRC1 and PRC2. We show that the depletion of sparse CBX2 leads to the dispersion of PRC1 and PRC2 condensates and results in the migration of PRC1 and PRC2 to constitutive heterochromatin, suggesting that sparse CBX2 is critical for proper spatial clustering of PRC1 and PRC2. Functionally, the depletion of CBX2 causes the migration of H3K27me3 to constitutive heterochromatin and the genome-wide reprogramming of H3K27me3. How do these results unify earlier observations to understand the facultative heterochromatinization of Polycomb target genes? It has been shown that the facultative heterochromatinization of Polycomb targets by PRC2 is through a two-step mechanism: PRC2 binds to a set of nucleation sites to methylate H3K27 and generate H3K27me3, which in turn stimulates PRC2 activity to spread H3K27me3 to distal regions ^13–19^. This spreading is facilitated by spatial clustering of Polycomb target genes ^18^. Mechanistically, increasing the concentration of PRC2 through CBX2 phase separation facilitates both nucleation and H3K27me3 spreading since a local high concentration can enhance reaction rates. Consistent with this hypothesis, our earlier studies have shown that LLPS speeds up the on-rate of the binding of CBX2 to chromatin ^37^, and a recent study has indicated that phase-separated condensates play a role in the clustering of Polycomb target genes ^96^. Taken together, these results suggest that sparse CBX2 nucleates PRC1 and PRC2 to assemble Polycomb condensates to create a high local concentration of PRC2 and to cluster Polycomb target genes, which facilitates H3K27me3 deposition and demarcates the spatial boundaries of H3K27me3 domains. Perturbation of CBX2 LLPS impairs facultative heterochromatinization of Polycomb target genes and cellular differentiation.

### Broad implications in biology

The discovery of the functional importance of sparse CBX2 has broad implications for biology. Many proteins are low in abundance, particularly proteins related to development since their expression is progressively turned on or off during development. In biology, it is widely held that low-abundant proteins are less functionally important, which is partly due to the fact that there are many limitations when studying low-abundant proteins, including their detection and quantification. Our findings challenge this preconception since we find that low-abundant proteins seed the formation of large assemblages by nucleating many other factors through phase separation. The assembled condensates can then generate isolated territories of enriched molecules, which enhance the efficiency and specificity of biochemical reactions. We expect that with the advent of ultra-sensitive single-molecule methods and genetic engineering, the understanding of many functionally relevant proteins with low abundance will accelerate.

## STAR METHODS

### RESOURCE AVAILABILITY

#### Lead contact

Further information and requests for resources and reagents should be directed to and will be fulfilled by the lead contact, Dr. Xiaojun Ren.

#### Materials availability

All stable/unique reagents generated in this study may be obtained from the lead contact with a completed material transfer agreement.

#### Data and code availability

- All data reported in this paper will be shared by the lead contact upon request.
- This paper does not report the original code.
- Any additional information required to reanalyze the data reported in this paper is available from the lead contact upon request.

### EXPERIMENTAL MODEL AND SUBJECT DETAILS

Unless otherwise indicated, female mouse embryonic stem cells (PGK12.1 provided by Dr. Neil Brockdoff from University of Oxford, UK) were cultured at 37°C and 5% CO_2_ on 0.2% gelatin, in mESC medium which consists of Dulbecco’s Modified Eagle Medium (DMEM) media supplemented with 15% Fetal Bovine Serum (VWR), 0.1 mM β-mercaptoethanol (Gibco), 2.0 mM L-Glutamine (Gibco), 100 Units/mL Penicillin-Streptomycin (Gibco), 1 × MEM Non-Essential Amino Acids (Gibco), 10 µg/mL Ciprofloxacin (Sigma), and 10^3^ units/mL leukemia inhibitor factor (LIF). To conditionally deplete CBX2, CBX2 ^fl/fl^-HaloTag mESCs were treated with 1.0 µM 4-Hydroxytamoxifen (Sigma) for 48 hours. To conditionally deplete fusion dTag Polycomb subunits, 100 nM dTag-13 (Tocris) was added to mESC medium for either 4 hours or 24 hours as noted.

For packaging used to develop the functional complementation mESCs, human HEK293T cells were cultured at 37 °C in 5% CO_2_ in MEF medium consisting of Dulbecco’s Modified Eagle Medium (Sigma) supplemented with 10% FBS, 0.1 mM β-mercaptoethanol, 2.0 mM L-Glutamine, 100 Units/mL Penicillin-Streptomycin, and 10 µg/mL Ciprofloxacin.

### METHOD DETAILS

#### Plasmid Generation

All Plasmids were generated in DH5α (Thermo Fisher Scientific).

##### HaloTag knockin donor plasmids for CRISPR/Cas9

To begin donor vector generation, additional restriction enzyme sites were added to the pUC19 vector. After the addition of extra restriction enzyme sites to pUC19, HaloTag vectors were prepared independently for both N- and C-terminal donor vectors. The N-terminal HaloTag insert was taken from FLAG-LoxP-PURO-LoxP-HaloTERT WT HR Donor (Addgene) which contains a 3 × FLAG epitope upstream of a puromycin resistance gene flanked by LoxP cassettes. The C-terminal HaloTag insert containing only HaloTag was amplified via PCR from the same vector. Both the N- and C-terminal HT inserts were cloned into the middle of the multiple cloning site of the modified pUC19 vector to give pUC19-N-HaloTag-LoxP-PURO and pUC19-HaloTag-C, respectively. An N-terminal HaloTag vector without LoxP sites or puromycin resistance was also generated via PCR to give pUC19-N-HaloTag. The right homology arm for the N-terminal HaloTag-CBX2 was amplified via PCR from mESC genomic DNA and the left homology arm was synthesized by Thermo Fisher. Ligation of the N-terminal CBX2 homology arms into the N-terminal pUC19-N-HaloTag-LoxP-PURO vector gave pUC19-N-HaloTag-CBX2. Both homology arms for the C-terminal CBX2-HaloTag donor vectors were amplified via PCR from mESC genomic DNA and ligated into pUC19-HaloTag-C to give pUC19-CBX2-HaloTag-C. The homology arms for the C-terminal RING1B-HaloTag donor vector were amplified via PCR from mESC genomic DNA and ligated into pUC19-HaloTag-C to give pUC19-RING1B-HaloTag. Similarly, C-terminal homology arms for CBX7, RING1A, MEL18, and EZH2 were amplified via PCR of mESC genomic DNA and added pUC19-HaloTag-C to give pUC19-CBX7-HaloTag, pUC19-RING1A-HaloTag, pUC19-Mel18-HaloTag, and pUC19-EZH2-HaloTag respectively. The N-terminal homology arms for PHC1 were amplified via PCR from mESC genomic DNA and added to pUC19-N-HaloTag to give pUC19-HaloTag-PHC1.

##### Conditional knockout donor plasmids for CRISPR/Cas9

To generate the conditional knockout pUC19-CBX2^fl/fl^-HaloTag donor vector, a region containing the last CBX2 exon, which comprises the majority of CBX2, and ∼200 bp of the upstream intron of was amplified via PCR from mESC genomic DNA. The PCR amplicon was then swapped with the original left homology arm from pUC19-CBX2-HaloTag-C. To add a LoxP site upstream of the left arm and to introduce base mutations compatible for the second sgRNA target site, a new DNA fragment containing a LoxP site and base mutations and spanning the last intronic region of *Cbx2* was synthesized by Thermo Fisher. The synthesized DNA fragment was added to the 5’ end of the insert from the previous step. A 3’ LoxP cassette was then added via oligo at the 3’ end of HaloTag and gave our pUC19-CBX2^fl/fl^-HaloTag donor vector. To insert ERT2CreERT2, an sgRNA was designed to increase integration efficiency of pROSA26-ERT2CreERT2 (Addgene) into the constitutively expressed ROSA26 locus.

##### dTAG-Venus donor plasmids for CRSIPS/Cas9

For the development of the dual knockin cells, a dTag-Venus insert was synthesized by Thermo Fisher containing a FKBP12F36V mutant and a gene for the Venus fluorescence protein. The synthesized insert was swapped with HaloTag in pUC19-HaloTag-C vector to give pUC19-dTag-Venus. To generate PHC1 dTAG-Venus, homology arms were amplified via PCR and cloned into the pUC19-dTag-Venus vector to give pUC19-dTag-Venus-PHC1. To generate CBX7 C-terminal Venus-dTag from, both homology arms were amplified via PCR and ligated into the original MCS-modified pUC19 vector. Then PCR amplified Venus was inserted first and PCR amplified dTag second to give pUC19-CBX7-Venus-dTag. Both homology arms for *Ring1b* and *Ezh2* were taken from their HaloTag plasmid counterparts and swapped with the *Cbx7* homology arms in pUC19-CBX7-Venus-dTag to give the pUC19-RING1B-Venus-dTag and pUC19-EZH2-Venus-dTag donor vectors, respectively.

##### Donor plasmids to generate a CBX2 mutant cell line by CRISPR/Ca9

For CBX2^DM^-HaloTag donor vector, an insert containing positive amino acids to alanine in the ATL and HPCR motifs was synthesized and swapped with the same region from the pUC19-CBX2^fl/fl^-HaloTag donor vector to generate the pUC19-CBX2-ATL-HPCR-P2A donor vector.

##### Guide RNAs for CRSPR/Cas9

The N-terminal HaloTag-CBX2 sgRNA oligos were inserted into pX330-U6-Chimeric_BB-CBh-hSpCas9 (Addgene). All other sgRNA oligos were cloned into pSpCas9(BB)-2A-Puro (PX459) V2.0 (Addgene). The guide RNAs were designed using the IDT’s ‘design custom gRNA’ tool, and sgRNA oligos synthesized from IDT were annealed.

##### Plasmids for protein expression

To generate plasmids for protein expression, the pGEX-6p-1 vector (Sigma) was used ^36^. YFP-FLAG was generated via PCR, ligated into pGEX-GST to generate the plasmid pGEX-GST-YFP-FLAG. Wild-type human CBX2 cDNA was amplified via PCR from pTRIPZ (M1)-HT-Cbx2 ^36^ and inserted to give pGEX-GST-CBX2-YFP-FLAG. To create the positive to alanine (P2A) mutations within the ATH, ATL, and HPCR motifs for the CBX2^TM^ protein expression vector, PCR primers were developed to change alanine residues, and individual fragments were assembled to give the complete vector. The PRC1 expression vectors containing RING1B, MEL18, and PHC1 without a fluorophore were generated in a previous study ^39^.

##### Plasmids for functional complementation

For the functional complementation assay, the pTRIPZ-YFP-CBX2 plasmid used in transduction has been previously described in ^36^. PCR was used to amplify the mutant insert from pGEX-GST-CBX2^TM^-YFP-FLAG, then the amplified insert was then swapped with *Cbx2* from the pTRIPZ-YFP-CBX2 vector to give pTRIPZ-YFP-CBX2^TM^.

#### Cell Line Generation

##### Generation of HaloTag knockin cell lines via CRISPR/Cas9

Approximately 2 million mESCs were mixed with 25 µg donor vector, 15 µg of sgRNA, and Ingenio electroporation buffer (Mirus Bio) to reach a total volume of 600 μL and incubated on ice for 10 minutes. Electroporation was performed using a Gene Pulser XCell (Bio-Rad) with the settings of 260V & 500 μF using an exponential decay in a 0.4 cm cuvette (Bio-Rad). After electroporation, cells were incubated on ice for an additional 10 minutes before being added to warm medium and plated on a gelatinized plate. For the cell lines generated with PX459 sgRNA, the morning after electroporation, 2.0 µg/ml puromycin (Sigma) was added to the cell medium for 24 hours, then reduced to 1.0 µg/ml puromycin for another 24 hours. Following puromycin selection for 2 days plates were washed twice with PBS before culture was resumed with mESC medium without puromycin. After approximately 1-week, colonies were picked and then expanded from 96-well plates to 24-well plates. A portion of each cell colony was split to an imaging dish and labeled with 100 nM HaloTag TMR ligand (Promega), and fluorescent colonies were identified using HILO microscopy. Genomic DNA from fluorescent colonies was purified, and homozygous or heterozygous insertion was verified via PCR. For the HaloTag-CBX2 cell line generated with PX330 sgRNA, since the donor vectors contain puromycin selection mark, cells were cultured with constant 1.0 µg/ml puromycin selection. Individual colonies were picked after a week and then genomic DNA was purified. Proper insertion was verified via PCR. Following PCR verification of the homozygous N-terminal tagging of HaloTag-CBX2, selected homozygous cell lines were transfected with 15 µg pBS598-EF1alpha-EGFPCre (Addgene) via electroporation and then were sorted with FACS for positive transient expression of GFP. The knockout was verified via PCR. Colonies that had either a homozygous or heterozygous insertion were further screened and validated with a western blot using Anti-HaloTag (Promega) and compared with the parental cell line as a negative control.

##### Generation of CBX2^fl/fl^-HaloTag conditional knockout via CRISPR/Cas9

To generate the endogenously tagged CBX2 conditional knockout cell line, referred to as CBX2^fl/fl^-HaloTag, Electroporation and puromycin selection were performed as before with 25 µg donor vector, 15 µg of each sgRNA vectors, and Ingenio electroporation buffer (Mirus Bio) to reach a total volume of 600 μL. Following insertion and validation of the proper HaloTag insert, pROSA26-ERT2CreERT2 (Addgene) was used as a donor vector along with another designed sgRNA to insert ERT2CreERT2 into the constitutively expressed ROSA26 locus. Cells were selected for one week with 6.5 µg/ml blasticidin (Gibco). Surviving colonies were picked, expanded, and imaged after 24 hours of addition of 1.0 µM 4-Hydroxytamoxifen (Sigma). Selected cells were then further expanded, their genomic DNA was purified, and PCR was run after knockout to conclude proper deletion of the expected region within the CBX2 genomic locus.

##### Generation of dual knockin cell lines via CRISPR/Cas9

To generate dual knock-in cell lines we used our CBX2^fl/fl^-HaloTag as a parental cell line. Electroporation and puromycin selection were performed as described above, except cells surviving selection were then taken for FACS and fluorescent cells were expanded. After expansion, individual colonies were generated via serial dilution and expanded. Genomic DNA was purified from these cell lines and homozygous or heterozygous knockin of the endogenously tagged loci were verified with PCR.

##### Generation of functional complementation cells

To generate cell lines that YFP-*Cbx*2 and YFP-*Cbx*2^TM^ were stably integrated into the genome of CBX2^fl/fl^-HaloTag mESCs, pTripZ-M1 plasmids containing either YFP-CBX2 or YFP-CBX2^TM^ were purified by using EndoFree Max Plasmid Kit (QIAGEN). HEK293T were seeded in a 100-mm dish the day before to reach no more than 85-95% confluency the day of transfection. 3-4 hours before transfection MEF medium was changed. 21 µg pTripZ, 21 µg psPAX2 (Addgene), and 10.5 µg pMD2.G (Addgene) were mixed with 130 µL 2.0 M CaCl_2_ (Sigma) and sterile Milli-Q water to give a total solution volume of 1050 µL. While slowly vortexing, 2×HBSS (50 mM HEPES pH 7.1 (Sigma), 280 mM NaCl, 750 µM NaHPO_4_, 750 µM NaH_2_PO_4_) was added the solution dropwise. The solution was then incubated on ice and monitored for no more than 10 minutes until it appeared slightly milky. As soon as slight precipitation was observed, the solution was distributed evenly dropwise over the plate of HEK293T cells. No more than 12 hours after transfection, the MEF medium was changed with 10 mL fresh culture medium, cells were incubated for 48 hours at 37°C, 5% CO_2_, and fusion of the cells was observed. Viral titer suspended in medium was collected and the cell debris was pelleted by centrifugation at 1,000 × g for 5 minutes at 4°C. A single-cell suspension of approximately 0.5 × 10^6^ CBX2^fl/fl^-HaloTag mESCs was obtained, transferred to a new 15 mL tube, and collected via centrifugation at 500 × g for 5 minutes at 4°C. The CBX2^fl/fl^-HaloTag cells were added to 10 mL virus-containing MEF medium 8 µg/mL polybrene (Sigma). Cells were plated on a gelatin-coated plate, incubated overnight, and 12-16 hours later the virus containing medium was removed, the plate was washed, and 10 mL fresh mES medium was added. After 24 hours the cells were washed once with PBS and fresh culture medium was added along with 2 µg/mL puromycin to select for cells that had proper integration of transduced genes. After one week of puromycin selection, cells were incubated for 24 hr with 0.1 µg/mL doxycycline hyclate (Sigma) and sorted by FACS to select for fluorescent colonies containing our stably integrated CBX2 genes. After selection cells were cultured with at least 1 µg/mL puromycin. Selected cells were further expanded and verified for proper integration via fluorescence microscopy.

#### Quantification of Polycomb Proteins within Cells

To determine the number of the Polycomb subunits within condensates, RING1B-HaloTag cells were chosen as the baseline because of their relative high abundance versus the other Polycomb subunits. Cells were split and a nuclear extraction was performed as outlined in the ‘***Nuclear extraction****’ section*. Three quantities of the RING1B-HaloTag nuclear extract from a known number of cells were ran against a serial dilution of HaloTag^®^ Standard Protein (Promega) and probed with Anti-HaloTag (monoclonal) (Promega) on a western blot as outlined in the ‘***Western Blotting***’ section. Intensities from each trial were quantified using the measurement tool in ImageJ and fit on the concentration curve generated by the serial dilution of the recombinant HaloTag^®^ Standard Protein so that we could generate a total number of RING1B proteins per cell. Following the quantification of RING1B-HaloTag per cell we quantified images of at least 10 cells from each of the endogenously tagged HaloTag fusion cell-lines under the same settings for their intensities and condensed fraction to be quantified as outlined in the ‘***CellProfiler analysis in vitro and live-cell condensate properties****’*. Total protein levels were generated ratiometrically using RING1B-HaloTag as the baseline versus average cell intensity ratios generated with CellProfiler. Standard deviation was calculated using the fluorescent intensities quantified. The number of proteins per condensate was calculated using the average number of proteins per cell, average number of condensates per cell, and the average condensed fraction.

#### Differentiation of mESCs to Embryoid Bodies and Neuron Progenitor Cells

##### EB differentiation

To differentiate mES cells to embryoid bodies (EBs), 2-5 x 10^6 cells were split and resuspended to single cells. Cells were then spun down at 200 × g for 5 minutes, washed with PBS, and spun again. After washing medium was replaced with 10 mL cell differentiation medium (CDM) which consists of Dulbecco’s Modified Eagle Medium supplemented with 15% Fetal Bovine Serum, 0.1 mM β-mercaptoethanol, 2.0 mM L-Glutamine, 100 Units/mL Penicillin-Streptomycin, 1 × MEM Non-Essential Amino Acids, 10 µg/mL Ciprofloxacin and plated onto a 100 mm petri dish (Falcon, 351029). For EB differentiation in the presence of retinoic acid (RA), 500 nM RA was added to CDM on day 4 of EB formation, and again on day 6. Every other day EBs were collected with a broken tip serological pipette, allowed to settle, and CDM was changed. EBs were collected at day 4 for NPC generation or continued until day 8 to be imaged or plated on a gelatin-coated plate for the EB spreading assay.

##### NPC differentiation

Before neuron progenitor cell (NPC) generation, polyornithine and laminin coated plates were prepared by adding 3 mL of 15 µg/mL polyornithine (Sigma) in Milli-Q water to sterile 60 mm cell culture dishes and left overnight at room temperature in a biological safety hood. The following day plates were washed 3 times with Milli-Q water and 3 mL of 1.0 µg/mL laminin (Sigma) in PBS was added, incubated at 37 °C overnight, and then frozen at −20 °C until use. The procedure and materials needed for differentiation from EBs to neuronal progenitor cells (NPC) has been previously described in ^45^. To further differentiate EBs to NPCs, EBs were collected on day 4 and transferred to the prepared polyornithine/laminin coated 60 mm plates with 5 mL of fresh CDM. After 24 Hours with CDM in coated plates, CDM was removed and changed to ITSFn medium which is DMEM/F12 supplemented with 5.0 µg/mL insulin (Sigma), 50 µg/mL apo-transferrin (Sigma), 30 nM selenium tetrachloride (Sigma), 5.0 µg/mL fibronectin (Sigma), 100 Units/mL Penicillin-Streptomycin, and 10 µg/mL Ciprofloxacin and cultured for an additional 6-8 days changing medium every other day to select for nestin-positive cells. After plating onto coated plates, images of outgrowth from EBs were taken with phase contrast imaging at time periods indicated. To expand nestin-positive cells, cells were gently but thoroughly dissociated with 0.05% trypsin, neutralized with CDM, spun down at 200 × g for 5 minutes, washed once with ITSFn medium, and spun down again. The ITSFn medium was then aspirated and replaced with 5 mL mN3FL medium which is DMEM/F12 supplemented with 25 µg/mL insulin (Sigma), 50 µg/mL apo-transferrin (Sigma), 30 nM selenium tetrachloride (Sigma), 20 nM progesterone (Sigma), 100 µM putrescine (Sigma), 1.0 µg/mL laminin (Sigma), 100 Units/mL Penicillin-Streptomycin (Gibco), and 10 µg/mL Ciprofloxacin (Sigma) and plated onto the new polyornithine/laminin coated plates. 25 ng/mL bFGF (R&D Systems) is added every day and mN3FL medium is changed every other day. For immunostaining and imaging, each NPC cell-type plate was dissociated with 0.05% trypsin, stopped with MEF medium, washed with PBS, and 1/3 of the cells were plated onto prepared polyornithine and laminin coated cover slides in a 6-well dish so that differences in growth could be observed. Cells were then fixed and stained as described in the “**Immunofluorescence**” section and imaged as outlined in the “***Optical setup for confocal microscopy***” section. Condensate properties and single-molecule dynamics were acquired with the same setup and settings as was used for the mESCs.

#### Preparation of Cellular Extracts

##### Nuclear extraction

To prepare nuclear extracts, cells were cultured on a p100 plate to 80-90% confluency to obtain ∼5-10 × 10^6 cells needed for extraction. Cells were first washed with PBS and detached from plate with 5 mL dilute sodium citrate solution (135 mM KCl and 15 mM sodium citrate in PBS) for 5 minutes. Detached cells were transferred to 5 mL mESC medium and collected at 200 × g for 10 minutes then washed once with ice-cold PBS and centrifuged again. PBS was removed and 7 mL hypotonic buffer (10 mM HEPES pH 7.5, 1.5 mM MgCl_2_, 10 mM KCl, and 0.1 mM PMSF) was gently added to the pellet and spun at 300 × g for 5 minutes. The buffer was removed, and the pellet was gently resuspended with 6 mL of fresh hypotonic buffer and incubated on ice for 10 min. Following incubation on ice the hypotonic solution containing the cells was added to a type B pestle then struck slowly and steadily 12 times to lyse the cells. Nuclei were collected by centrifugation at 1000 × g for 3 minutes, hypotonic buffer was removed, and 200 µL of nuclear lysis buffer (20 mM Tris-HCl pH 7.4, 2% NP-40 (IGEPAL^®^ CA-630), 300 mM NaCl, 0.25 mM EDTA, 20% Glycerol, 0.1 mM Na_3_VO_4_, 0.5 mM DTT, 1 × Protease Inhibitor Cocktail (Sigma), and 0.1 mM PMSF) was added. Cell nuclei were incubated at 4 °C for 30-60 minutes until lysis was complete. Afterwards nuclear lysis debris was collected by centrifugation at max speed and supernatant was collected. Before long-term storage at - 20°C, sample concentrations were quantified on a Qubit (Thermo scientific) so concentrations could be normalized before western blotting and 100 µL of 4x Laemmli Sample Buffer (Bio-Rad) & 30 µL 100 mM DTT (Sigma) were added to the nuclear extracts.

##### Histone extraction

To prepare histone extracts relevant cell lines were cultured on a p100 plate to 80-90% confluency to obtain ∼5-10 × 10^6 cells needed for extraction. Cells were first washed with PBS and detached from plate with 5 mL dilute sodium citrate solution (135 mM KCl and 15 mM sodium citrate in PBS) for 5 minutes. 5 mL mES medium was added to cells and collected at 200 × rcf for 10 minutes at 4°C. Cells were then resuspended with 1 mL ice-cold PBS, transferred to 1.5 mL Eppendorf tube, and centrifuged at 200 × rcf for 10 minutes at 4°C. PBS was removed gently as to not disturb the pellet and cells were resuspended with 1.0 mL of ice-cold hypotonic buffer (10 mM HEPES pH 7.5, 1.5 mM MgCl_2_, 10 mM KCl, 0.25 mM EDTA, mM Na_3_VO_4_, 0.5 mM DTT, 1 × Protease Inhibitor Cocktail (Sigma), and 0.1 mM PMSF. After addition of hypotonic buffer, cells were incubated on ice for 15 minutes before adding 50 µL of 10% NP-40 (IGEPAL^®^ CA-630) and vortexed for 10 seconds at max speed. Nuclei were then collected by centrifugation at 2,000 × rcf for 10 minutes at 4 °C and supernatant was removed. To lyse the nuclei 400 µL of 0.2 M H_2_SO_4_ was added, the pellet was resuspended completely, and the tube was rocked at 4 °C for a least 1 hour or overnight. After nuclear lysis the solution containing the histones was centrifuged at 10,000 × rcf for 10 minutes at 4 °C to remove debris. To precipitate histones the supernatant was then transferred to a new 1.5 mL Eppendorf tube and 200 µL TCA (Sigma) was added 4 drops at a time and tubes were inverted several times to mix between additions until the final TCA concentration reached 33%. Following precipitation, the samples were incubated for 30 minutes or overnight on ice and histones were collected by centrifugation at 16,000 × rcf with a soft ramp for 20 minutes at 4°C. Supernatant was then carefully removed and the histone pellets were washed with 1 mL ice-cold acetone and collected by centrifugation at 16,000 × rcf with a soft ramp for 10 minutes at 4°C twice. After the second acetone wash, supernatant was removed down to 5-10 µL and histone pellets were left to air dry for 5 minutes to remove residual acetone. Pellets were then dissolved in 200 µl of Milli-Q water with 100 mM Tris-HCl pH 7.4 & 200 mM NaCl. Before long-term storage at −20°C, sample concentrations were quantified on a Qubit (Thermo Scientific) so concentrations could be normalized before western blotting and 100 µL of 4x Laemmli Sample Buffer (Bio-Rad) & 30 µL 100 mM DTT was added to the histone extracts.

#### Western Blotting

Nuclear and histone extracts premixed with Laemmli Sample Buffer and DTT were denatured by heating to 95 °C for 10 minutes. Samples were then first diluted or directly loaded and resolved in a NuPAGE™ 4-12% Bis-Tris Gel (Thermo Scientific) and ran with the setting of 400 mAmp & 110V (constant) for 60-140 minutes. Before transfer to a nitrocellulose membrane, extra thick blotter paper is soaked in transfer buffer (48 mM Tris-base, 39 mM glycine, 0.037% SDS, and 20% methanol in Milli-Q water) and placed on a Trans-Blot^®^ SD Semi-Dry Transfer Cell (Bio-Rad) and bubbles are carefully removed with a roller. Then an Immobilon®-FL PVDF Membrane (Millipore) is soaked in pure methanol for 15 seconds before being placed on the blotter paper and removing bubbles. After placement of the nitrocellulose membrane, the Bis-Tris Gel containing the proteins is carefully placed on top of the membrane, bubbles are removed with a roller, and then another piece of extra thick blotter paper is added to the stack and the last bubbles are removed. The semi-dry cell was then reassembled, and proteins were transferred to the membrane with the setting of 20 V and 200 mAmps for 40 minutes. After transfer the membrane is placed in blocking buffer, which is comprised of TBS-T (20 mM Trizma-base, 136 mM NaCl, 3.8 mM HCl, and 0.1% Tween-20 in pH 7.6 Milli-Q water) and 5% dry milk for at least 2 hours. Prior to the addition of the primary antibody, the membrane is washed 3 times with TBS-T for 20 minutes. After blocking and washing the membrane was incubated with primary antibodies diluted in TBS-T with 1% dry milk overnight as follows: anti-HaloTag (monoclonal) (1:2000, Promega); anti-H3K27Me3 (1:10000, Cell Signaling); anti-RING1B (1:5000, MBL); anti-EZH2 (1:2000, Millipore). The following day membranes were washed 3 times with TBS-T Buffer for 20 minutes and then incubated at room temperature for 2 hours with either anti-Rabbit IgG Horseradish Peroxidase (1:5000, Azure) or anti-Mouse IgG Horseradish Peroxidase (1:5000, GE Healthcare) corresponding to the primary antibody host. After incubation with the secondary antibody, membranes were again washed 3 times with TBS-T for 20 minutes and the proteins were visualized using Radiance Plus chemiluminescent HRP substrate (Azure) with an Azure 300 imager (Azure). Following visualization membrane was stripped for 40 minutes with shaking using NewBlot™ PVDF Stripping Buffer (LI-COR) and then blocked with blocking buffer. After blocking again, membranes were incubated for 2 hours with either anti-GAPDH (FL-335) rabbit polyclonal antibody (1:4000; Santa Cruz) for the nuclear extracts or anti-H3 (1:5000, Cell Signaling) for the histone extracts, then washed as before, and incubated for 2 hours with anti-Rabbit IgG Horseradish Peroxidase linked whole antibody (1:5000, Azure). Following 3, 20-minute washes with TBS-T the membranes were visualized as described above for the controls.

#### Recombinant Protein Purification

The expression generated pGEX-6P-1-GST-FLAG vectors containing CBX2, CBX2^DM^, CBX2^TM^ with a fluorophore, and RING1B, MEL18, or PHC1 without a fluorophore were transformed into Rosetta™ 2 (pLysS) host strains (Novagen). A single colony was inoculated in 5.5 mL LB broth with 100 µg/mL ampicillin at 37°C for 16-18 hours while shaking at 250 rpm. The following morning, 4.0 mL of overnight culture was transferred to 1.0 L of fresh LB and shook at 250 rpm, 37°C for 7-8 hours to reach OD600 ≥ 1.5. To induce protein expression, 1.0 mM IPTG was added to the culture and the culture was shaken at 18°C for 16-18 hours. E. Coli was harvested via centrifugation at 5,000 × g for 15 minutes at 4°C and resuspended in 25 mL lysis buffer (50 mM HEPES pH 7.5, 0.5 mM EDTA, 1.6 M KCl, 0.5 mM MgCl_2_, 1 mg/mL lysozyme, 1 mM DTT, 0.2 mM PMSF, 20 µg/mL RNase A, and 1 protease inhibitor tablet (Sigma)). Three freeze-thaws cycles, alternating between −80°C and an ice-water bath were used to partially lyse bacteria. Cells were completely lysed via sonication with a Vibra-Cell™ (Sonics) for 7.0 minutes using 65% amplitude and cycles at 15 seconds on-time, 45 seconds off-time (28 cycles total). NP-40 (IGEPAL^®^ CA-630) was added to 0.1% and the solution was rocked at 4°C for 30 minutes before centrifugation at 15,000 × g for 15 minutes to pellet cell debris. Cell debris is discarded, supernatant is transferred to new tube, and DNA and RNA is precipitated with 0.30% PEI which was added dropwise while solution was carefully vortexed. After nucleic acid precipitation, the solution was rocked for 10 minutes at 4°C before DNA/RNA was collected by centrifugation at 20,000 × g for 10 minutes at 4°C. To ensure complete DNA/RNA precipitation, the same volume of PEI was added again dropwise to a concentration 0.30% and the solution was rocked for another 20 minutes at 4°C. Remaining DNA/RNA was collected by centrifugation at 20,000 × g for 10 minutes. Supernatant was added to 750 µL Pierce™ Glutathione Agarose (Thermo Scientific) pre-washed with cold 1× PBS and rocked for 1 hour at 4°C while protected from light. Beads were collected via centrifugation at 700 × g for 2 minutes at 4°C and supernatant was carefully and completely removed. Beads with bound protein were washed with 8-10 mL chilled Buffer A (20 mM HEPES pH 7.5, 0.2 mM EDTA, 1.0 M KCl, 0.2 mM PMSF, and 1 mM DTT) and collected by centrifugation at 700 × g at 4°C for 2 minutes. Buffer A from wash was removed and beads with bound protein were transferred to a Poly-Prep Chromatography Column (Bio-Rad) where they were washed twice with 10 mL chilled Buffer A and allowed to drain by gravity. To elute recombinant protein from Pierce agarose beads, 500 µL of GSH Elution Buffer pH 8.0 (Buffer A + 80 mM reduced L-Glutathione + Tris-Base to pH) was incubated for 10 minutes at 4°C and then collected. Elution was repeated with 3 × 500 µL total additions to reach a 1.5 mL elution volume. Collected protein was incubated 1 hour with 200 µL ANTI-FLAG-M2 affinity resin (Sigma) pre-washed with 3 times with 1.0 mL Buffer B (20 mM HEPES pH 7.5, 0.2 mM EDTA, 1.0 M KCl, 0.2 mM PMSF, and 1 mM DTT). Affinity resin was collected via centrifugation at 500 × g at 4°C for 2 minutes. Resin was washed a total of three times with 1.0 mL Buffer B and collected by centrifugation to remove supernatant. To elute protein, ANTI-FLAG resin was rocked with 300 µL Buffer BC (20 mM HEPES pH 7.5, 0.2 mM EDTA, 0.05% NP-40, 20% glycerol, 0.5 mM DTT, 0.1 mM PMSF, 1 × protease inhibitor cocktail (Sigma), 300 mM KCl, and 0.8 mg/mL FLAG peptide (Sigma)) at 4°C for 1 hour. Resin was collected via centrifugation at 2,000 × g for 2 minutes at 4°C. Supernatant was then collected and transferred to a new 1.5 mL tube before being spun at 20,000 × g for 10 minutes to remove all remaining affinity resin. The supernatant was collected and transferred to a 10K MWCO 0.5 mL Pierce™ concentrator (Thermo Scientific) where it was concentrated down to 20-30 µL by centrifugation at 15,000 × g for 45-50 minutes at 4°C in a fixed-angle rotor. Following the purification of recombinant RING1B, MEL18, and PHC1, the GST epitope was removed with Precision Protease according to the manufacturer’s recommendations. Recombinant protein was resolved by NuPAGE™ 4-12% Bis-Tris Gel (Invitrogen) to determine identity and purity. Protein concentration was quantified by measuring absorption at 595 nm with a Nanodrop^TM^ 2000 (Thermo Scientific) using the Pierce™ Detergent Compatible Bradford Assay Reagent (Thermo Scientific) against serial BSA standards.

#### Labeling of Recombinant PRC2 with Alexa Fluor^®^ 555

Before the *in vitro* condensate formation assay, recombinant PRC2 (Active Motif) was labeled with Alexa Fluor^®^ 555 using a protein labeling kit (Thermo Scientific) and a vial from an Antibody Labeling Kit (Thermo Scientific). 1 M sodium bicarbonate was prepared by adding 1 mL of Milli-Q to provided vial in Antibody Labeling Kit (Thermo Scientific) and vortexed until fully dissolved. A dilution buffer (25 mM HEPES pH 7.5, 300 mM NaCl, 0.04% Triton-X100 in Milli-Q) was prepared and added to the recombinant PRC2 complex in a 1:1 ratio to a total volume of 80 µL. 10 µL (1/10th volume of protein solution to be labeled) of 1 M sodium bicarbonate was added for a final volume of 90 µL PRC2 complex solution. 0.5 mL Dilution Buffer was added to the included Alexa Fluor^®^ 555 from the kit, and 10 µL of the diluted Alexa Fluor^®^ 555 to 90 µL of diluted PRC2 complex with 0.1M sodium bicarbonate and gently inverted multiple times to mix thoroughly. Solution was incubated for 1 hr at room temperature and inverted every 10-15 minutes to increase labeling efficiency. While the dye solution was incubating, a spin-column for the removal of excess dye from labeled proteins was prepared by ensuring 2 frits were inserted at the bottom of the spin column and adding 1.0 mL premixed purification resin and allowing for settling by gravity. More purification resin was added to a total bed volume of 1.5 mL and buffer was drained from column by gravity after centrifuging for 3 minutes at 1100 x g. Buffer exchange was performed on resin in column by adding 500 µL of Dilution Buffer 3 times. After 1 hr dye incubation, 100 µL reaction mixture was added to prepared resin column with new collection tube and centrifuged for 5 min at 1100 x g. Glycerol was added to 5% and DTT was added to 0.5 mM to final solution before long-term storage at −80°C.

#### *In Vitro* Condensate Formation Assay

For the individual CBX2 condensate formation assay serial dilutions of the purified GST-YFP-CBX2-FLAG, GST-CBX2^DM^-YFP-FLAG, and GST-CBX2^TM^-YFP-FLAG in were made using additional BC Buffer (20 mM HEPES pH 7.5, 0.2 mM EDTA, 0.05% NP-40, 20% glycerol, 0.5 mM DTT, 0.1 mM PMSF, 1 × protease inhibitor cocktail (sigma), 300 mM KCl, and 0.8 mg/mL FLAG peptide (Sigma) to the following concentrations: 10 µM, 5 µM, 2.5 µM, 1.0 µM, 0.5 µM, 0.1 µM, 0.05 µM, and 0.01 µM. After dilution in BC Buffer, samples were then further diluted 1:10 with ice-cold imaging buffer (20 mM HEPES pH 7.5, 1 mM MgCl_2_, and 70 mM KCl in Milli-Q water) to give a final concentration of 100 mM KCl. After 1:10 dilution with imaging buffer and gentle mixing, samples were left to sit at room temperature for 20 minutes before being added to a perfusion chamber CoverWell™ Perfusion Chamber (Grace Bio-Labs) adhered to a sonicated and MeOH cleaned 25 × 25 mm micro cover glass (VWR). Holes on the perfusion chamber were taped to prevent evaporation and condensates were allowed to settle for around 5 minutes before acquiring ∼20 images for each condition. For the condensate formation assay with recombinant CBX2, labeled PRC2 complex, Cy5-Nucleosome, and PRC1 components RING1B, MEL18, and PHC1 recombinant protein samples were diluted with BC buffer to concentrations of 5 µM, 1 µM, 2 µM, 5 µM, 5 µM, and 5 µM respectively. After dilution in BC Buffer, samples were then further diluted 1:10 together with ice-cold imaging buffers such that the final buffer concentration is 20 mM HEPES pH 7.5, 1 mM MgCl_2_, and 100 mM KCl. After thorough mixing diluted recombinant proteins with the proper imaging buffers to give the necessary salt concentrations, samples were incubated as before. Separate assays for CBX2 only, PRC2 only, CBX2+PRC2, CBX2+Nucleosome, PRC2+Nucleosome, and CBX2+PRC2+Nucleosome were performed and ∼15-20 images were taken with the same settings for each wavelength. Similarly, a separate series of assays was performed for CBX2+PRC2+Nucleosome+RING1B, CBX2+PRC2+Nucleosome+RING1B+MEL18, and CBX2+PRC2+Nucleosome+RING1B+MEL18+PHC1. Images were acquired as described in the “***Optical setup for in vitro condensate imaging***” section and analyzed to give a condensed fraction as described in the “***CellProfiler analysis of in vitro and live-cell condensate properties***” section.

#### Immunofluorescence

For each of the immunofluorescence experiments, approximately 0.5 × 10^6 cells were seeded in each well of 6-well plate containing a gelatinated cover glass and incubated overnight at 37°C, 5% CO_2_. The following day, the cells were washed with PBS and fixed with 1% formaldehyde for 10 minutes at room temperature. Cells were then washed with PBS and permeabilized by incubating with 0.2% Triton X-100 for 10 minutes with gentle shaking and washed again twice with PBS. The cells were incubated with blocking buffer (10.0 mM PBS pH 7.4, 0.1% Triton X-100, 3% Goat Serum, 3% w/v BSA) overnight. After washing with basic blocking buffer (10.0 mM PBS pH 7.4, 0.1% Triton X-100), the primary antibody was diluted in blocking buffer as follows: anti-H3K27Me3 (1:4000, Cell Signaling); anti-nestin (1:10, DSHB, rat-401-S). Diluted primary antibodies were then added to washed cells and incubated for 2 hours at room temperature. After incubation with primary antibody, cells were washed with basic blocking buffer and secondary antibodies were diluted in blocking buffer as follows: Alexa Fluor 488-Goat anti-Mouse IgG (H+L) secondary antibody (1:1000, Life Technologies) or Alexa Fluor 568-Goat anti-Rabbit IgG (H+L) secondary antibody (1:1000, Life Technologies). After washing, cells were incubated with diluted secondary antibodies and covered for 2 hours at room temperature. After incubation with the secondary antibody, cells were washed and incubated with 0.1 µg/mL Hoechst stain at room temperature while covered for 10 minutes. After a final wash with PBS, the cover glass was mounted to a frosted microscope slide using ProLong^®^ Gold Antifade reagent (Invitrogen) and left flat to dry overnight. Images of DAPI/Hoechst and the other fluorescent protein of interest were acquired using the setup described in “***Optical setup for confocal microscopy***” section and the same settings were used across various conditions used in quantification. Images were analyzed for enrichment as outlined in the “***ImageJ analysis for protein enrichment versus hoechst***” section. Images were processed for presentation by Photoshop and ImageJ.

#### RNA Extraction, qPCR, and RNA-Seq

RNA was extracted from mESC and EBs using an RNEasy Kit (QIAGEN) according to the manufacturer’s recommendations and filtered pipette tips. After extraction samples were quantified with Qubit (Thermo Scientific). RT was performed using the ProtoScriptII cDNA Synthesis Kit (NEB) to give cDNA. For qPCR, cDNA from each sample was ran in triplicate with primers designed to amplify 180-220 bp spanning exons of each target gene of interest on a StepOne™ Real-Time PCR System (Applied Biosystems), and GAPDH was ran in triplicate for each sample to normalize expression. For mRNA sequencing of wild-type and *Cbx2* KO (with OHT for two days) mESCs, 1.0 µg of total RNA per sample was used to generate sequencing libraries with NEBNext® Ultra™ II Directional RNA Library Prep Kit for Illumina®. following manufacturer’s instructions. The library was then quantified by using Qubit dsDNA HS Assay kit and examined using a TapeStation High Sensitivity DNA assay kit. The prepared mRNA library was then sequenced on Nextseq 500 to generate ∼ 20 million PE75 reads.

#### CUT&RUN

The CUT&RUN of CBX2 in CBX2^fl/fl^-HaloTag was performed using a CUTANA™ ChIC/CUT&RUN Kit (EpiCypher). The knockout of CBX2 induced by the addition of OHT for 48 hours was used as a negative control. Cells were harvested, split to a single-cell suspension, counted, and 0.5 x 10^6 cells were used in each reaction, and reactions were run in duplicate. Cells were washed with PBS, then Wash buffer (20 mM HEPES pH 7.5, 150 mM NaCl, 0.5 mM Spermidine, 1 x EDTA-free protease Inhibitor), and bound to activated ConA beads for 10 minutes. Supernatant was removed from beads, 50 μL Antibody Buffer ((20 mM HEPES pH 7.5, 150 mM NaCl, 0.5 mM Spermidine, 1 x EDTA-free protease Inhibitor, 0.1% Digitonin, 2 mM EDTA) with 1:50 Anti-HaloTag^®^ (polyclonal) (Promega) or 1:200 Anti-GFP (monoclonal) (Invitrogen) was added, and cells were gently vortexed to thoroughly resuspend beads before overnight incubation at 4°C on a nutator. The following day, the Antibody buffer was removed with the tubes on a magnetic stand and cells were washed twice with pAG-MNase Buffer (20 mM HEPES pH 7.5, 150 mM NaCl, 0.5 mM Spermidine, 1 x EDTA-free protease Inhibitor, 0.1% Digitonin). After the washes, 50 μL pAG-MNase Buffer + 2.5 μL pAG-MNase was added to each reaction, tubes were then vortexed to mix and incubated at RT for 10 minutes. Supernatant was removed and each reaction was washed twice with 200 μL pAG-MNase Buffer. Afterwards 2 mM CaCl_2_ in pAG-MNase Buffer was added to activate pAG-MNase for chromatin digestion, beads were vortexed to resuspend, and tubes were then incubated at 4°C on a nutator for 2 hours. Stop buffer (340 mM NaCl, 20 mM EDTA, 4 mM EGTA, 50 μg/mL RNase A, 50 μg/mL Glycogen) + E. coli spike-in was added following digestion and tubes were gently vortexed to mixed then placed in a preheated thermocycler set to 37°C. Reactions were then purified and eluted from included columns, then quantified using a Qubit. Following quantification samples were library prepped using a NEBNext^®^ Ultra™ II DNA Library Prep Kit for Illumina^®^. Following end repair and adaptor ligation, 1X SPRIselect beads were used to wash prepped DNA and PCR was ran for 15 cycles of denaturation and annealing/extension. The PCR settings were: Initial Denaturation at 98°C for 30 Seconds, Denaturation at 98°C for 10 seconds, Annealing/extension at 65°C for 75 seconds, Final Extension at 65°C for 5 minutes. Following PCR, 1X SPRIselect beads were used to wash amplified library-prepped DNA and each eluted product was ran on a Tapestation (Agilent) for fragment analysis before multiplexing and PE150 sequencing to a depth of at least 8 million reads. Data was analyzed as outlined in the ‘**NGS Analysis**’ section.

#### CUT&Tag

CUT&Tag was performed as described before ^97^. Genome-wide H3K27me3 domain changes were profiled using the CUT&Tag protocol adapted from EpiCypher. Cells were harvested via 0.25% trypsin-EDTA solution and counted using 0.4% trypan blue stain and Countess automated cell counter. A single-cell suspension was prepared by washing cells with cold 1x PBS and resuspending 1.0 x 10^6^ cells in 1 mL cold nuclear extraction buffer (20 mM HEPES-KOH pH 7.9, 10 mM KCl, 0.1% Triton x-100, 20% glycerol, & 0.5 mM spermidine), then incubated on ice for 10 minutes to isolate nuclei. Biomag Plus Concanavalin A beads were activated with two washes of cold bead activation buffer (20 mM HEPES-KOH pH 7.9, 10 mM KCl, 1 mM CaCl_2_, & 1 mM MnCl2_2_). After activation, beads were added to aliquots of 0.1 x 10^6^ isolated nuclei and incubated at room temperature for 10 minutes. Nuclei:bead samples were resuspended in 50 µL cold Antibody150 buffer (20 mM HEPES-KOH pH 7.5, 150 mM NaCl, 0.5 mM spermidine, 0.01% digitonin, 2 mM EDTA, & protease inhibitors) followed by the addition of 0.5 µg of anti-H3K27me3 (#9733S, Cell Signaling Technology). To facilitate primary antibody binding, samples were incubated on nutator at room temperature for 1 hour. Samples were resuspended in 50 µL Digitonin150 buffer (20 mM HEPES-KOH pH 7.5, 150 mM NaCl, 0.5 mM spermidine, 0.01% digitonin, & protease inhibitors) followed by the addition of 0.5 µg CUTANA anti-rabbit secondary antibody (#13-1047, EpiCypher). Samples were then incubated on nutator at room temperature for 1 hour to facilitate binding, followed by two washes with 200 µL cold Digitonin300 buffer (20 mM HEPES-KOH pH 7.5, 300 mM NaCl, 0.5 mM spermidine, 0.01% digitonin, & protease inhibitors). The tagmentation reaction was performed by resuspending samples in 50 µL cold Tagmentation buffer (20 mM HEPES pH 7.5, 300 mM NaCl, 0.5 mM spermidine, 10 mM MgCl_2_, & protease inhibitors) and incubating at 37°C in thermocycler for 1 hour. Samples were washed with 50 µL TAPS buffer (10 mM TAPS pH 8.5 & 0.2 mM EDTA) then resuspended in 5 µL SDS Release buffer (10 mM TAPS pH 8.5 & 0.1% SDS) to quench the tagmentation reaction and release adaptor-tagged DNA fragments into solution. After incubating samples at 58°C in thermocycler for 1 hour, 15 µL SDS Quench buffer (0.67% Triton x-100), 2 µL i5 universal primer (EpiCypher), 2 µL i7 barcoded primer (EpiCypher), 0.005 ng *Escherichia coli* spike-in DNA (made in-house), and 25 µL NEBNExt High Fidelity 2x PCR Master Mix were added to each sample for library construction with the following PCR protocol: 5 minutes at 58°C, 5 minutes at 72°C, 45 seconds at 98°C, 14 cycles of 15 seconds at 98°C and 10 seconds at 60°C, with a final extension for 1 minute at 72°C. Libraries were checked for quality using the Qubit dsDNA HS Assay kit and TapeStation High Sensitivity DNA assay kit and then sequencing was performed using Novaseq 6000 (Novogene) to generate 150 bp paired-end reads. Data was analyzed as outlined in the ‘**NGS Analysis**’ section.

#### Optical Settings

##### Optical setup for confocal microscopy

Confocal images were captured using a Zeiss LSM 700 Microscope (Zeiss, Germany). For imaging of prepared immunofluorescence mES cells, an Alpha Plan-Apochromatic 100×/1.40 NA Oil-immersion Objective (Zeiss, Germany) was used. For imaging of prepared immunofluorescence NPC cells, an Alpha Plan-Apochromatic 20×/1.40 NA Oil-immersion Objective (Zeiss, Germany) was used. The microscope and the camera were controlled by Zen Black software (Zeiss, Germany).

##### Optical setup for in vitro condensate imaging

For the recombinant condensate formation assays, epifluorescence images were obtained using a Zeiss Axio Observer D1 Manual Microscopy (Zeiss, Germany) equipped with an Alpha Plan-Apochromatic 100×/1.46 NA Oil-immersion Objective (Zeiss, Germany) and an Evolve 512 × 512 EMCCD camera with pixel size 16 µm (Photometrics, Tucson, AZ). For the excitation and emission of YFP, a Brightline® single-band laser filter set (Semrock; excitation filter, FF02-482/18-25; emission filter, FF01-525/25-25; dichroic mirror, Di02-R488-25) was used. For the excitation and emission of Alexa 555, a Brightline^®^ single-band laser filter set (Semrock; excitation filter, FF01-561/14; emission filter, FF01-609/54; dichroic mirror, Di02-R561-25) was used. For Cy5 fluorescence the emission filter: FF01–640/14–25, and dichroic mirror: Di02-R635–25 3 36) was used. The microscope and the EMCCD camera were controlled by Slidebook 6.0 software.

##### Optical setup for live-cell condensate imaging

For imaging of condensates in living cells, images were obtained using a Zeiss Axio Observer D1 Manual Microscopy (Zeiss, Germany) equipped with an Alpha Plan-Apochromatic 100×/1.46 NA Oil-immersion Objective (Zeiss, Germany) was used. To avoid stray-light reflection and reduce background from cell auto-fluorescence, the HILO illumination model was used as described previously ^98^. A Solid-state LaserStack (3i) was used to excite TMR HaloTag® Ligand (Promega) and Venus, For the excitation and emission of TMR, a Brightline® single-band laser filter set (Semrock; excitation filter, FF01-561/14; emission filter, FF01-609/54; dichroic mirror, Di02-R561-25) was used. For the excitation and emission of Venus, a Brightline® single-band laser filter set (Semrock; excitation filter, FF02-482/18-25; emission filter, FF01-525/25-25; dichroic mirror, Di02-R488-25) was used. The microscope and the EMCCD camera were controlled by Slidebook 6.0 software (Intelligent Imaging Innovations, Colorado).

##### Optical setup for live-cell single-molecule tracking

For live-cell single-molecule tracking, an Axio Observer D1 Microscope equipped with an Alpha Plan-Apochromat 100 × /1.46 Oil immersion Objective (Zeiss, Germany) and an Evolve 512 × 512 EMCCD camera with pixel size 16.0 μm was used, along with an additional 2.5 × magnification on the emission pathway. A Solid-state LaserStack (3i) was used to excite Janelia Fluor® 549 HaloTag® Ligand at 552 nm. To avoid stray-light reflection and reduce background from cell auto-fluorescence, the HILO illumination model was used as described previously ^98^. A Brightline® single-band laser filter set (Semrock; excitation filter: FF01–561/14, emission filter: FF01–609/54, and dichroic mirror: Di02-R561–25 3 36) was used for the excitation and emission spectra of JF549. To filter the excitation wavelength, a TIRF laser microscope cube (3i) was used. The microscope and EMCCD camera were controlled by the computer via SlideBook 6.0 software. The images for population were acquired with the following settings in SlideBook 6.0: Intensification 700, Number of frames 100, Dark time (ms) 0, Exposure time (ms) 30, Laser power (mW) 7.5, TIRF angle ∼6.80. The images for residence time were acquired with the following settings in SlideBook 6.0: Intensification 700, Number of frames 400, Dark time (ms) 170, Exposure time (ms) 30, Laser power (mW) 2.25, TIRF angle ∼6.80.

#### Imaging Dish Preparation

Plates for live-cell imaging are first prepared by drilling a ½” hole into a 35 mm cell culture dish. The hole is cleaned of plastic debris with a razor and then the plates are washed with water to remove any additional plastic residue from drilling. After the plates dry, a sonicated and flame cleaned 22 × 22 mm cover slip is attached to the bottom of the plate with a small bead of 5-minute epoxy (Devcon) around the cleaned hole so that the slip sits flush with the dish and left to dry for at least 24 hours. After sterilization with 70% ethanol for 10 minutes, a wash with PBS, and UV in a biological safety cabinet for at least 15 minutes, plates are gelatinated and incubated at 37 °C for at least 2 hours.

#### Live-cell Imaging of Condensates

Approximately 0.5 × 10^6 cells were seeded in each imaging dish and incubated overnight. To image cells with HaloTag fusion proteins, 50 nM TMR dye (Promega) was added to cells for independent or dual fluorescent imaging and incubated at 37 °C for 15 minutes. Dye was then washed off with PBS, mES medium is added, and cells were placed back into the incubator for 30 minutes so that cells could recover. After an additional PBS wash and 30-minute recovery with mES medium, the plate was washed again with PBS and then imaging medium (FluoroBrite DMEM (Life Technologies)) supplemented with 10% FBS and 10^3^ units/mL LIF) was added before imaging. Cells were imaged for 60-90 minutes while temperature was held at 37 °C using a Single Channel Temperature Controller (Warner Instruments). Images were acquired using the configuration listed in “***Optical setup for live-cell condensate imaging***” section. Images were analyzed for number of condensates, condensed fraction, and intensity as described in the “***CellProfiler analysis of in vitro and live-cell condensate properties***” section. Images were prepared for presentation using Photoshop and ImageJ.

#### Live-cell Single-molecule Tracking

Approximately 0.5 × 10^6 cells were seeded in each plate for imaging and incubated overnight. For labeling of our mES or NPC cells with HaloTag fusion proteins, 50-500 pM Janelia Fluor® 549 (Tocris, 6147) dye was added to cells in culture medium such that 5-20 proteins are visible within each nucleus. Kinetic and chromatin bound population dynamics were analyzed as previously described ^24,37,99,100^. Images were acquired using the configuration listed in “***Optical setup for live-Cell single-molecule tracking***” section.

#### Single-molecule Photobleaching to Quantify CBX2 within Condensates

CBX2^fl/fl^-HaloTag cells were plated to ∼20% confluency on imaging dishes prepared with 0.2% gelatin and cultured overnight. The following day cells were incubated with Janelia Fluor^®^ 549 HaloTag^®^ Ligand at 50 pM & 100 pM concentrations for the sparsely labeled condition and 500 nM for the fully labelled condition. After incubation with dye for 15 minutes, cells were protected from light and washed gently with PBS followed by a 30-minute recovery with mESC medium in the incubator. A wash and recovery were performed twice before fixation. After recovery cells were washed twice with PBS and then fixed for 10 minutes with 1% formaldehyde directly on the imaging dishes. After fixation cells were washed once with PBS, then imaged in PBS. HILO videos of cells were acquired in with a bright time of 30 ms and a dark time of 0 ms. Low laser intensities were used such that bleaching events would occur over the span of 2-4 minutes until complete bleaching allowing for more precise quantification. Foci were outlined in ImageJ with the ROI Manager and time traces of fluorescent intensity were generated from the image stacks. The photobleaching steps were detected and quantified after applying Chung-Kennedy filtration ^101^ to the image stack intensities. Each photobleaching step corresponds to one protein within each focus. Data was acquired from 8-15 cells and a total of 150-300 points were quantified per condition.

#### Image Analysis

##### ImageJ analysis for protein enrichment versus hoechst

Cells were analyzed for enrichment of the protein of interest versus Hoechst with the following pipeline: cells for analysis were identified, background intensity proximal to cell on image corresponding to Hoechst stain was selected with ‘Polygon selections’ tool and imported to the ‘ROI Manager’; cell on image corresponding to Hoechst stain was outlined with ‘Polygon selections’ tool and imported to the ‘ROI Manager’; all high-intensity regions within cell on image corresponding to Hoechst stained dense heterochromatin were outlined with ‘Polygon selections’ tool and imported to the ‘ROI Manager’ independently; measure function used to extract intensity and area data; intensity and area data corresponding to Hoechst stain was exported to spreadsheet; ROI for Hoechst intensity was normalized versus background; image corresponding to protein of interest was selected on ImageJ; regions of interest saved in the ‘ROI Manager’ were highlighted on the image corresponding to protein of interest (H3K27me3/Venus); intensity and area data corresponding to protein of interest was extracted with the measure function; intensity and area data corresponding to protein of interest was exported to spreadsheet; intensity was normalized vs background; enrichment was determined by variation of normalized (intensity*area)/total(intensity*area) per cell between the Hoechst stain image and the corresponding protein of interest image; enrichment ratio of each selected region from each cell was exported to a main spreadsheet to determine standard deviation within the groups and p-value was calculated using Student’s t-Test between day 0-2 and 0-4 days of the knockout timed series. 150+ ROIs were analyzed for enrichment from each condition/time-point.

##### CellProfiler analysis of in vitro and live-cell condensate properties

For the quantification of condensates *in vitro*, two sets of images were taken under the same conditions for the quantification of *in vitro* condensates: one image was taken of buffer only for measurement of background fluorescence used in normalization, and the second image was of the buffer and each protein at various concentrations specified. We used the following pipeline to identify and quantify condensates formed by the recombinant proteins: identify primary object, measure object intensity, measure image intensity, export data to spreadsheet. The following parameters were used to identify primary objects: object size in pixel units, 5-40; upper outlier fraction, 0.05; averaging method, mean; variance method, standard deviation; # of deviations, 20; threshold smoothing scale, 1.35; threshold correction factor, 1.0; and lower and upper bounds on threshold, 0 – 1.0. The condensed fraction was calculated as follows:

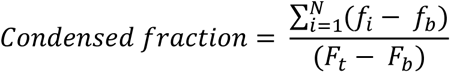

where *f*_*i*_ is the fluorescence intensity of individual condensates, *f*_*b*_is the background fluorescence intensity of individual condensates, *F*_*t*_ is the total fluorescence intensity of condensed and diluted parts, and *F*_*b*_ is the total background fluorescence intensity.

Condensate size was analyzed as described previously ^39^. The cumulative frequency distributions of condensate sizes in pixel were fitted with a lognormal cumulative distribution function based on the F-test implemented in OriginLab with:

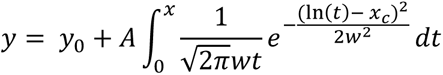

where the size of condensates is *x*_*c*_.

For the quantification of condensates in living cells we used the following pipeline to first identify then measure condensates: crop cell of interest, crop region without cells for background fluorescence baseline, enhance or suppress features, identify primary objects, measure image intensity, measure object intensity, and export data to spreadsheet. To identify primary objects the following settings were used: object size in pixel units 2 - 20; threshold strategy, adaptive; threshold method, robust background; lower outlier fraction, 0.05; upper outlier fraction, 0.05; averaging method, mean; variance method, standard deviation; # of deviations, 1.01; threshold smoothing scale, 0; threshold correction factor, 1.02; lower and upper bounds on threshold, 0 - 1.0; and size of adaptive window, 10. The condensed fraction and condensate size were described as above for the *in vitro* condensates. At least 10 cells/images were analyzed for each fluorescent protein of interest in both *in vitro* and live-cell experiments.

##### Live-cell single-molecule tracking analysis

Kinetic and chromatin bound population dynamics were analyzed as previously described ^24,37,99,100^. For residence time and kinetic fraction analysis, image stacks without visible cell drift were analyzed. Tracking settings used in U-track algorithms ^102^ were applied in MATLAB as described previously ^24,99,100^. To determine kinetic fractions, we constructed the displacement histograms and then carried out kinetic modeling of the measured displacements using Spot-On ^103^, which quantitatively measures three kinetic fractions (populations) of total molecules within the nucleus. The defined populations are F_1_ (total chromatin bound), F_2_ (confined motion), and F_3_ (free diffusion). All Spot-On settings were described previously ^24,37,99,100^. To determine stable chromatin-bound fraction (F_s_) and long-residence time (τ_l_), we determined the track length of individual particles whose *D*_*m*_ is less than 0.032 μm^2^/s. Following correction for photobleaching, the normalized cumulative frequency distributions are fitted with a one- or two-component exponential decay function as described previously ^24,37,99,100^, which generate stable chromatin-bound fraction (F_s_) and long-residence time (τ_l_). The target-search time (τ_s_) was estimated from the long residence time (τ_*l*_) and stable chromatin-bound fraction (*F_s_*) as described previously ^24,37^.

#### NGS Data Analysis

Data analysis was performed as previously described ^97^. CUT&Tag and CUT&RUN sequencing reads were trimmed using Cutadapt ^104^: Illumina adapter sequences were removed, and reads were trimmed to 140 bp. Reads less than 35 bp were discarded. Trimmed reads were aligned to the mm10 version of *Mus musculus* genome using bowtie2 ^105^. Samtools ^106^ and bedtools ^107^ were used for processing aligned reads from sam to bed files. Duplicate reads were discarded for further analysis if the reads had the same start and end coordinates. Coverage at 100 bp windows genome-wide was calculated as the number of reads that mapped at that window, normalized by the factor N:

N = 2800000000/(Total number of mapped reads)

2800000000 was a number chosen arbitrarily in the range of total mapped base pairs in M. musculus genome. The normalized read counts were then smoothed with a running average spanning +/- 1000 bp around each 100 bp bin. The distribution of normalized read counts in 100 bp windows genome-wide was generated, and a “domain cutoff” was determined as the normalized read count that is greater than the normalized read count of 95% of the windows with coverage. Domains were called by linking adjacent windows with a normalized read count ≥ domain cutoff. To account for short disruptions due to mappability issues, jumps of up to 750 bp were allowed while linking windows. A log2 ratio of H3K27me3 enrichment over IgG enrichment was calculated for all the putative domains. Those domains with a log2 enrichment greater than 2 (4-fold enrichment over IgG) were used for all downstream analyses. For performing disjoin and reduce operations, the GenomicRanges package in R was used ^93^.

For RNA-seq analysis, demultiplexed fastq files were trimmed using Trim Galore and were aligned to mm10 genome reference using STAR with default parameters. The mapping files were then counted using featureCounts to generate the raw count matrix across samples. The differential gene expression analysis was performed using DESeq2 package in R/Bioconductor. Gene ontology analysis was performed using ClusterProfiler package in R/Bioconductor.

## ACKNOWLEDGMENTS

We acknowledge the Ren lab past and current members for helping experiments during this project and stimulating discussion during the writing process. This work was supported by the NIGMS (National Institute of General Medical Sciences) under Award Number R01GM135286 (to X.R.), R35GM133434 (to S.R.), the RNA Bioscience Initiative, the University of Colorado School of Medicine (to S.R.), and the Office of Research Service at the University of Colorado Denver (Grand Challenge Funding).

## AUTHOR CONTRIBUTIONS

X.R. conceived and designed the study, supervised the experiments, analyzed data, prepared the figures, and wrote the paper. S.R. conceived and designed the experiments of CUT&Tag, performed bioinformatic analysis, prepared the figures, and wrote the paper. T.W. conceived and designed experiments of RNA sequencing and contributed to the writing of the manuscript. T.K. and C.P. interpreted the results and contributed to the writing of the manuscript. S.I., A.T., X.L, A.E., J.W., J.T., and K.A. performed experiments, analyzed data, and wrote the manuscript.

---

**Supplementary Figure 1.**
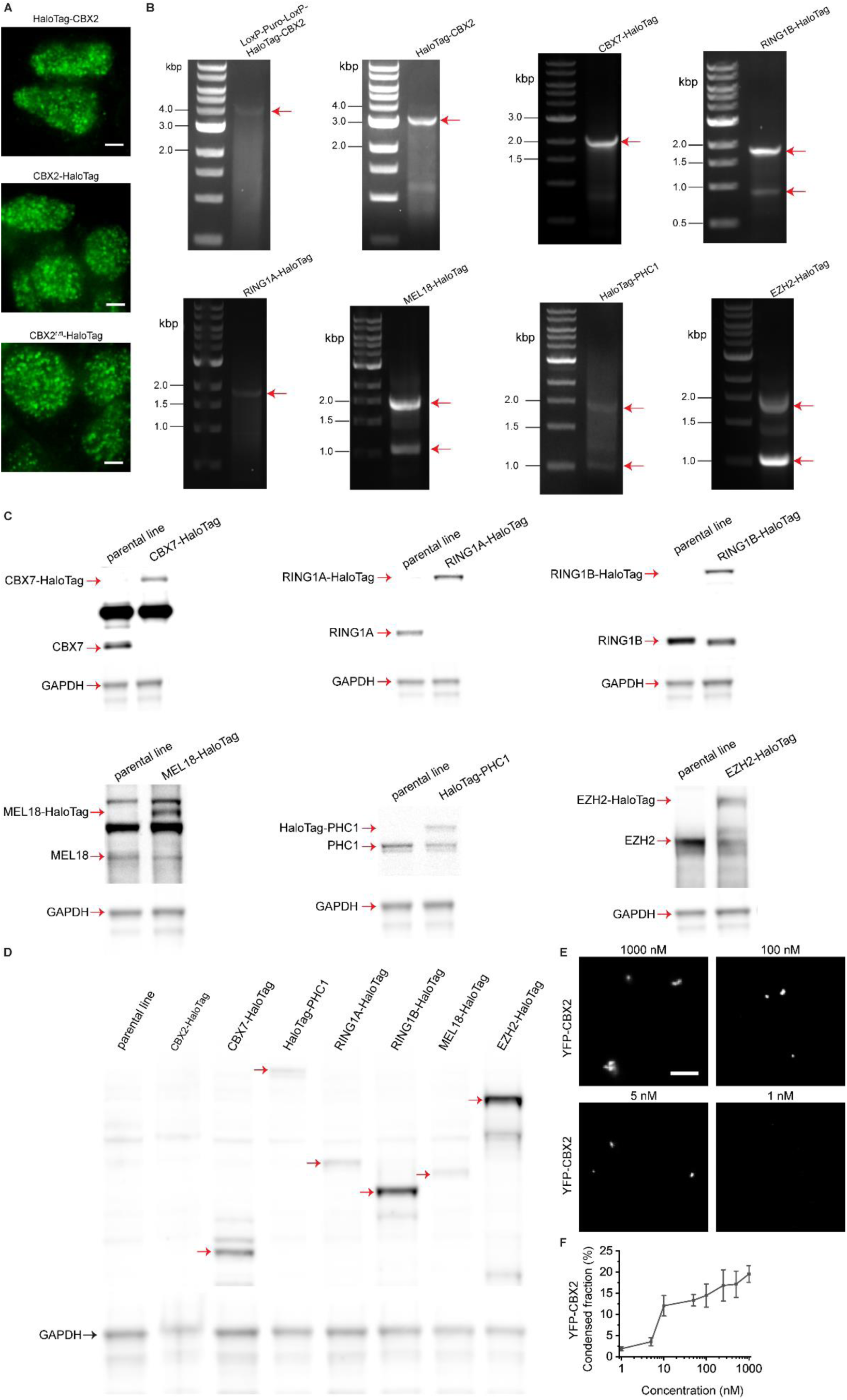
∼3 CBX2 proteins per condensate in mESCs. (**A**) Live-cell epifluorescence micrographs of HaloTag knockined at the N-terminus of *Cbx2* (HaloTag-CBX2) or at the C-terminus of *Cbx2* that is flanked without (CBX2-HaloTag) or with two LoxP sites (CBX2^fl/fl^-HaloTag). Images were taken by using single-molecule sensitive HILO microscopy. Scale bar, 5.0 µm. (**B**) Agarose gel analysis of PCR amplicons from mESCs where HaloTag was inserted into the endogenous loci of Polycomb genes. PCR primers span the junction of right- and left arms and the HaloTag insert. Red arrowheads indicate the correct band size of a predicted length. A single arrowhead indicates a homozygous knockin, and two arrowheads show a heterozygous knockin. (**C**) Immunoblotting analysis of HaloTag Polycomb fusion proteins isolated from their corresponding knockin mESCs as well as control parental mESCs. Gels were detected by antibodies against individual Polycomb proteins. Red arrowheads indicate the correct bands of fusion and endogenous proteins, respectively. GAPDH was used for control. (**D**) Immunoblotting analysis of HaloTag Polycomb fusion proteins isolated from their corresponding knockin mESCs and control parental mESCs. Gels were probed by using anti-HaloTag antibodies. Red arrowheads indicate the correct band of fusion Polycomb proteins. CBX2-HaloTag was too low to detect. GAPDH was used for control. Scale bar, 5.0 µm. (**E**) Representative epifluorescence images of condensates formed by YFP-CBX2 at concentrations of 1 nM, 5 nM, 100 nM, & 1000 nM. Condensates were imaged and processed by using the same settings. Scale bar, 5.0 µm. (**F**) Condensed fraction of CBX2 under a series of CBX2 concentrations.

**Supplementary Figure 2.**
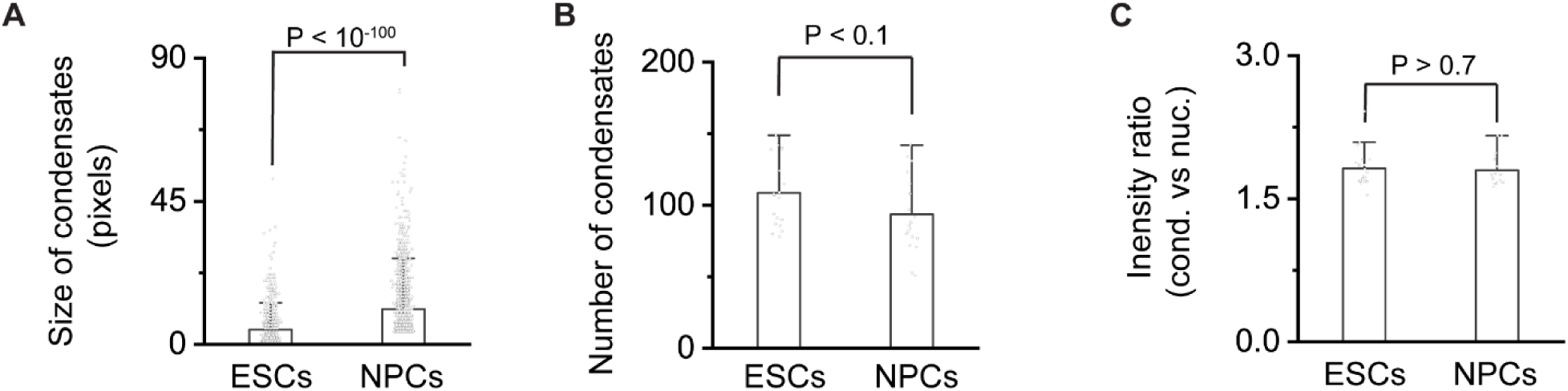
Cell-type-dependent control of the number of CBX2 proteins per condensate and the engaging of CBX2 with chromatin. (**A**) Bar overlap plot of the condensate size of CBX2-HaloTag in mESCs and NPCs. P value, student’s t-test with two-tailed distribution and two-sample unequal variance. (**B**) Bar overlap plot of the number of condensates of CBX2-HaloTag in mESCs and NPCs. P value, student’s t-test with two-tailed distribution and two-sample unequal variance. (**C**) Bar overlap plot of the fluorescence intensity ratio of condensates versus whole nucleus for CBX2-HaloTag in mESCs and NPCs. P value, student’s t-test with two-tailed distribution and two-sample unequal variance.

**Suppl Figure 3.**
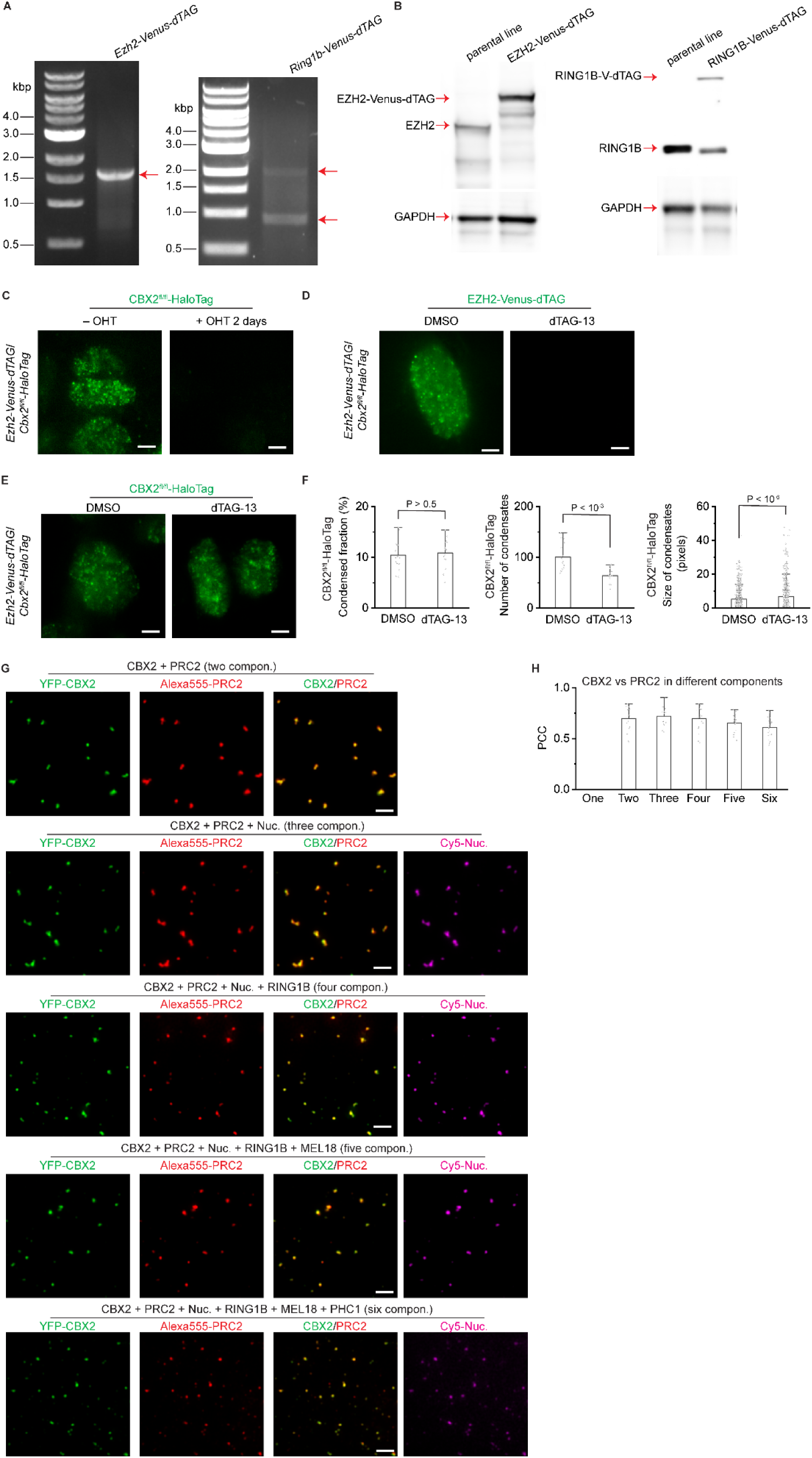
Sparse CBX2 nucleates Polycomb condensates. (**A**) Agarose gel electrophoresis analysis of PCR amplicons from *Cbx2^fl/fl^-HaloTag* mESCs where Venus-dTAG has been inserted into the C-terminus of *Ezh2* and *Ring1b* genes, respectively. Red arrowheads indicate the correct band of predicted size. One arrowhead represents homozygous knockin and two show heterozygous knockin. (**B**) Immunoblotting analysis of EZH2-Veus-dTAG and RING1B-Venus-dTAG isolated from their corresponding knockin mESCs as well as control parental mESCs. Gels were detected by antibodies against EZH2 and RING1B, respectively. Red arrowheads indicate the correct bands of fusion and endogenous proteins, respectively. GAPDH was used for control. (**C**) Live-cell epifluorescence images of CBX2-HaloTag in *Ezh2-Venus-dTAG*/*Cbx2^fl/fl^-HaloTag* mESCs that were treated with or without OHT for two days. Scale bar, 5.0 µm. (**D**) Live-cell epifluorescence images of EZH2-Veus-dTAG in *Ezh2-Venus-dTAG*/*Cbx2^fl/fl^-HaloTag* mESCs that were treated with dTAG-13 or DMSO for 24 hours. Scale bar, 5.0 µm. (**E**) Live-cell epifluorescence images of CBX2-HaloTag in *Ezh2-Venus-dTAG*/*Cbx2^fl/fl^-HaloTag* mESCs that were treated with dTAG-13 or DMSO for 24 hours. Scale bar, 5.0 µm. (**F**) Bar overlap plot of the condensed fraction, the number of condensates, and the size of condensates of CBX2-HaloTag in *Ezh2-Venus-dTAG*/*Cbx2^fl/fl^-HaloTag* mESCs that were treated with dTAG-13 or DMSO for 24 hours. P value, student’s t-test with two-tailed distribution and two-sample unequal variance. (**G**) Epifluorescence images of condensates of YFP-CBX2, Alexa555-PRC2, and Cy5-nucleosomes in heterogenous systems with different components. Individual and overlay images of CBX2 and PRC2 were shown. Images of nucleosomes were also shown. Scale bar, 5.0 µm. (**H**) Bar overlap plot of condensate-based Pearson correlation coefficient (PCC) of CBX2 and PRC2 quantified from Figure S3G.

**Suppl Figure 4.**
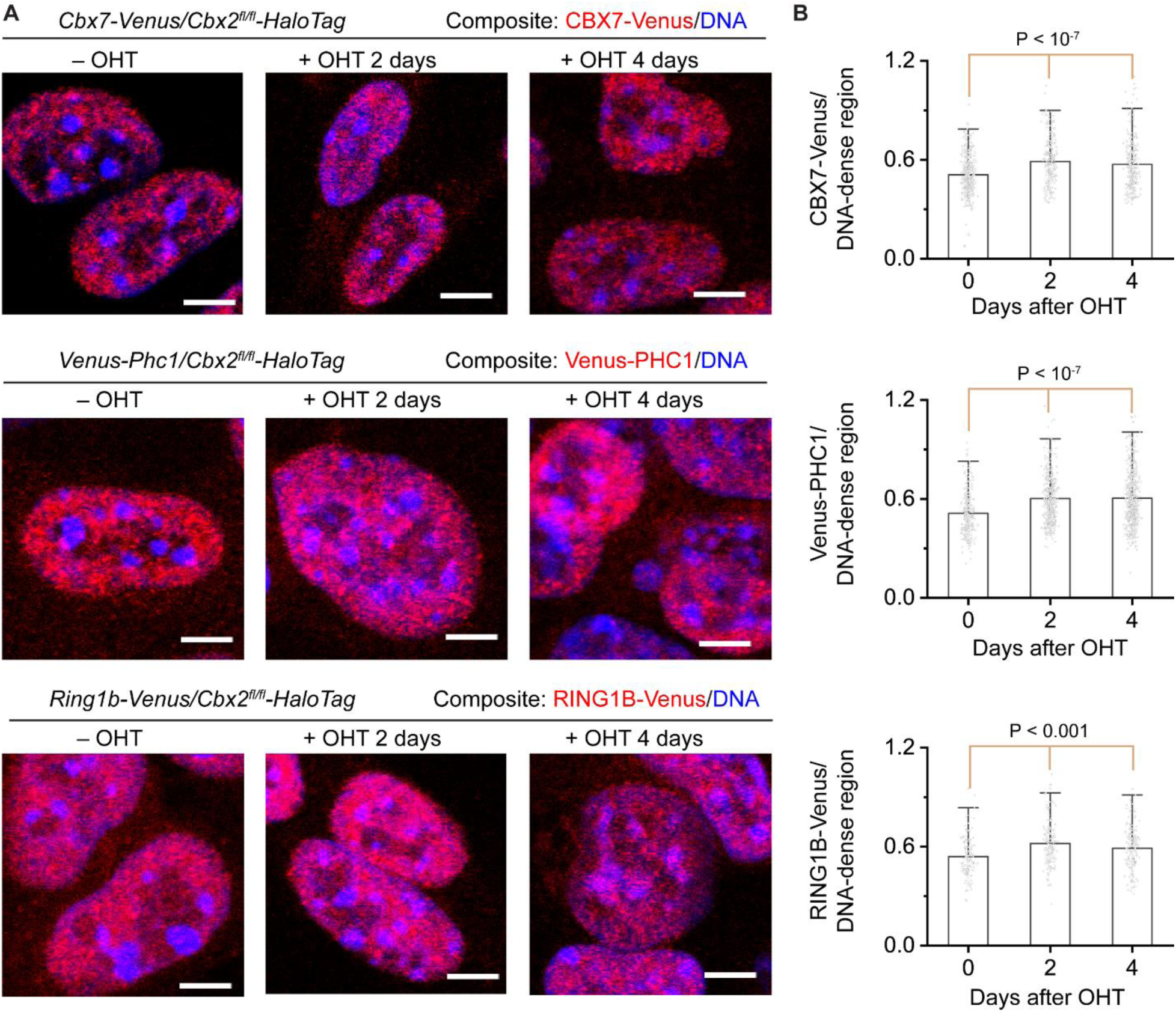
Sparse CBX2 demarcates the spatial boundaries of facultative heterochromatin and controls the H3K27me3 level. (**A**) Representative confocal images of CBX7-Venus, Venus-PHC1, and RING1B-Venus in *Cbx7-Venus/Cbx2^fl/fl^-HaloTag*, *Venus-Phc1/Cbx2^fl/fl^-HaloTag*, and *Ring1b-Venus/Cbx2^fl/fl^-HaloTag* mESCs, respectively, treated with or without OHT for different time periods. DNA was stained by Hoechst. Overlay images are shown. PRC1 subunit fusions are red, and DNA is blue. Scale bar, 5.0 µm. (**B**) Bar overlap plot of the fluorescence intensity ratio of CBX7-Venus, Venus-PHC1, and RNIG1B-venus to DNA dense regions quantified from Figure S4A. P value, student’s t-test with two-tailed distribution and two-sample unequal variance.

**Supplementary Figure 5.**
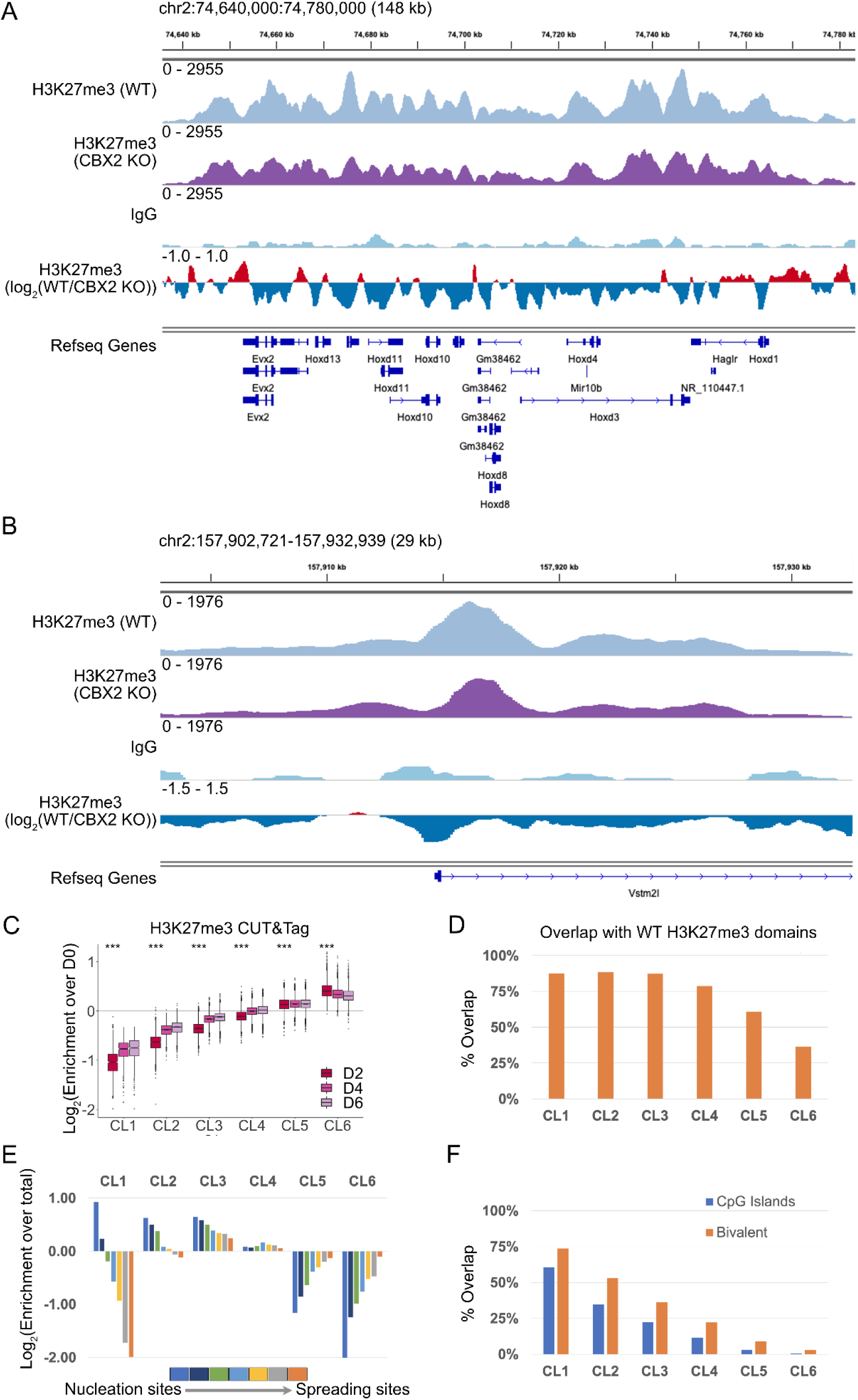
Effect of CBX2 loss on H3K27me3 domains. (**A**) Genome browser snapshot showing CUT&Tag enrichment for H3K27me3 in WT, CBX2 KO, and IgG, and log2 ratio of H3K27me3 enrichment in CBX2 KO over WT are shown for the Hoxd locus. (**B**) Same as (A) for a representative bivalent promoter. (**C**) Distribution of log2 fold change of H3K27me3 CUT&Tag enrichment for CBX2 KO over WT at segments defined in Figure 5A for each cluster shown as a boxplot. Wilcox rank sum test was used to calculate the p-value with the null hypothesis that the log2 fold change for each cluster on day 2 (D2) is 0, and p-values were adjusted for multiple comparisons using the Bonferroni correction. *** denotes p < 2.2e-16 (**D**) Percentage of segments from each cluster that overlap with H3K27me3 domains defined in WT. (**E**) Overlap of segments from each cluster with kinetic clusters that denote nucleation and spreading sites as defined in Veronezi et al ^19^. The log2 ratio of the overlap for each cluster over the overlap of all segments combined is plotted. (**F**) Percentage of segments from each cluster that overlap with CpG islands and bivalent promoters.

**Suppl Figure 6.**
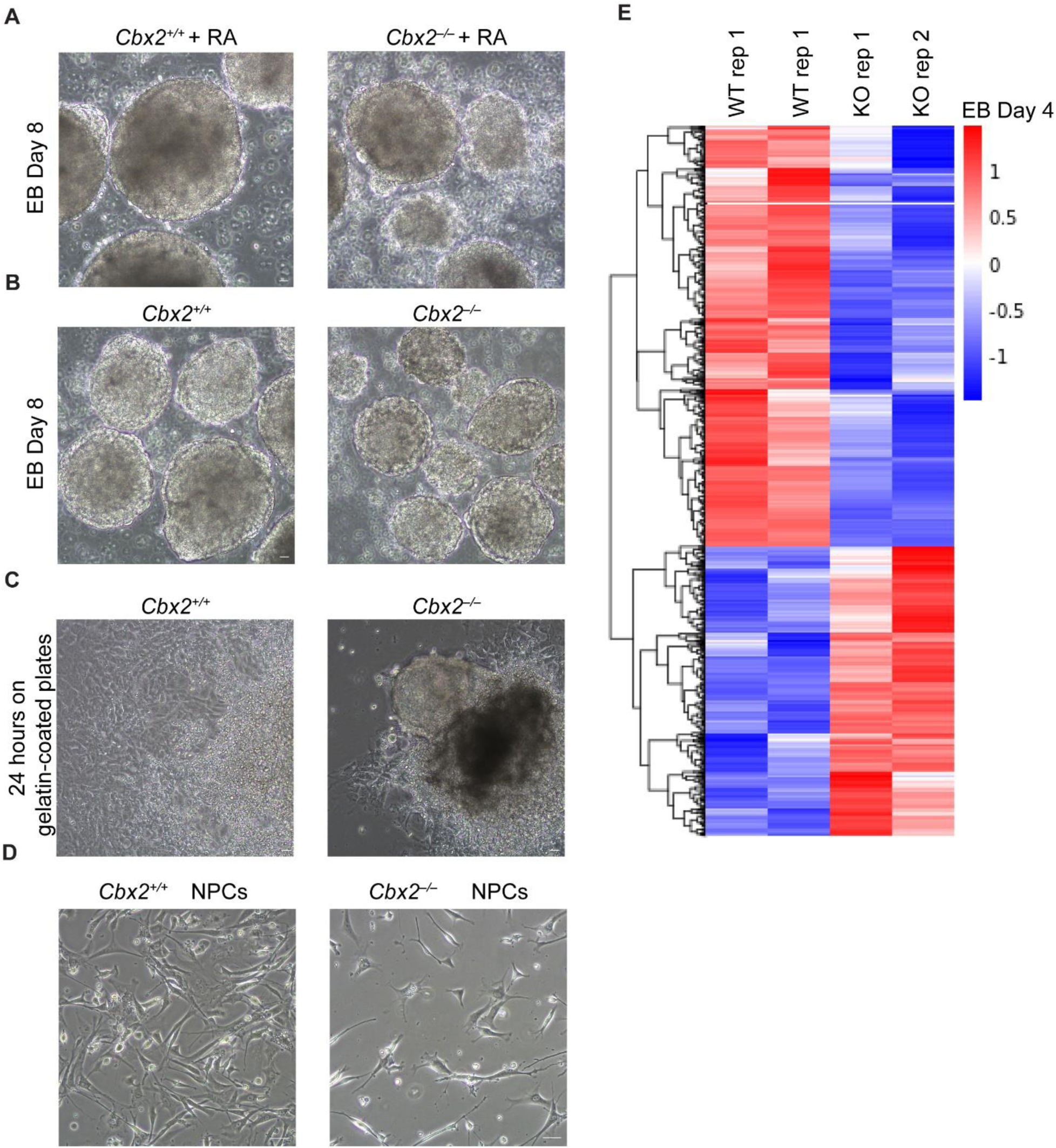
CBX2 is needed for cellular differentiation. (**A**) Phase-contrast images of day-8 EBs from wild-type (*CBX2^+^/^+^*) and knockout (*CBX2^−/−^*) cells in the presence of retinoic acid (RA). Scale bar, 100 µm. (**B)** Phase-contrast images of day-8 EBs from wild-type (*CBX2^+^/^+^*) and knockout (*CBX2^−/−^*) cells in the absence of retinoic acid (RA). Scale bar, 100 µm. (**C**) Phase-contrast images of the outgrowth of day-8 EBs from wild-type (*CBX2^+^/^+^*) and knockout (*CBX2^−/−^*) cells on gelatin-coated plates 24 hours after plating in the absence of retinoic acid (RA). Scale bar, 100 µm. (**D**) Live-cell phase-contrast images of NPCs derived from wild-type (*CBX2^+^/^+^*) and knockout (*CBX2^−/−^*) cells under the same differentiation conditions. Scale bar, 100 µm. (**E**) The hierarchical clustering analysis of the DEGs between wild-type (*CBX2^+^/^+^*) and knockout (*CBX2^−/−^*) from day-4 EBs. RNA expression values (RPKM, reads per kilobase per million reads) are represented by z-score across samples.

**Suppl Figure 7.**
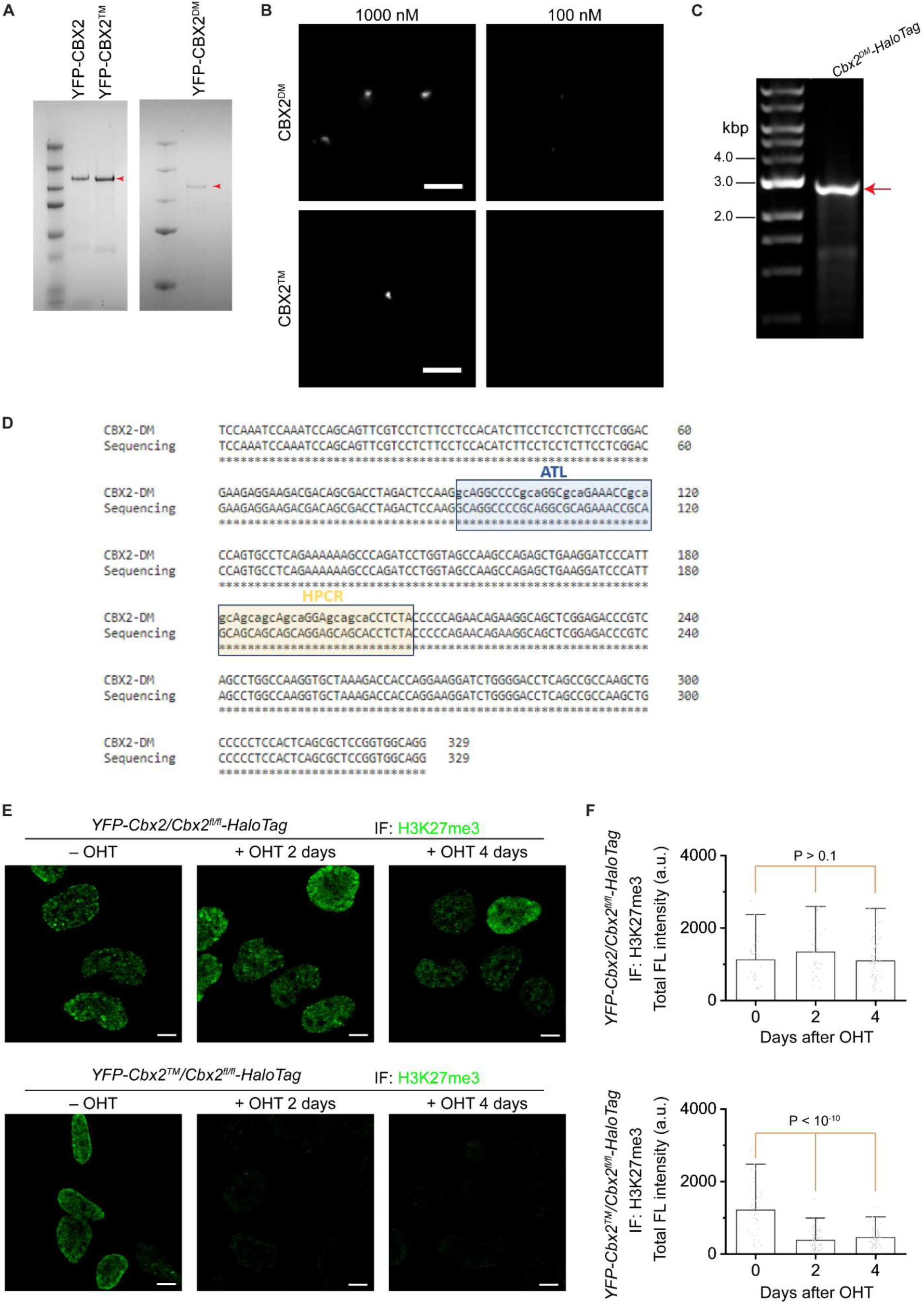
LLPS of CBX2 demarcates the spatial boundaries of facultative heterochromatin and regulates cellular differentiation. (**A**) SDS-PAGE analysis of recombinant YFP-CBX2, YFP-CBX2TM, and YFP-CBX2DM purified from E. coli. Red arrowheads indicate the correct bands of predicted molecular weight. (**B**) Representative epifluorescence images of condensates formed by YFP-CBX2DM and YFP-CBX2TM. Scale bar, 5.0 µm. (**C**) Agarose gel analysis of PCR amplicons from Cbx2DM-HaloTag mESCs. PCR primers span junction between the right- and left homology arms and HaloTag insert. Red arrowheads indicate the correct band size of predicted length. (**D**) Sanger sequencing analysis of PCR amplicons from Cbx2DM-HaloTag mESCs. ATL and HPCR are boxed. Lower-case letters are mutated bases. (**E**) Example epifluorescence images of immunostained H3K27me3 in YFP-Cbx2/Cbx2fl/fl-HaloTag and YFP-Cbx2TM/Cbx2fl/fl-HaloTag mESCs, respectively. Cbx2 was depleted by administrating OHT for 2 days. Scale bar, 5.0 µm. (**F**) Box overlap plot of the fluorescence intensity of immunostained H3K27me3 in YFP-Cbx2/Cbx2fl/fl-HaloTag and YFP-Cbx2TM/Cbx2fl/fl-HaloTag mESCs quantified from Figure S7E. P value, student’s t-test with two-tailed distribution and two-sample unequal variance.

